# The Disease Gene THAP12 is a Transcriptional Regulator of Mitochondrial ETC Complex I

**DOI:** 10.64898/2026.07.16.738975

**Authors:** Brandon R. Desousa, Yohei Abe, Martin Kampmann, Isha H. Jain

**Affiliations:** Gladstone Institute of Cardiovascular Disease, Gladstone Institutes, San Francisco, CA 94158, USA; Biomedical Sciences Graduate Program, University of California, San Francisco, San Francisco, CA 94158, USA; Arc Institute, Palo Alto, CA 94304, USA; Department of Biochemistry and Biophysics, University of California, San Francisco, San Francisco, CA 94158, USA; Institute for Neurodegenerative Diseases, University of California, San Francisco, San Francisco, CA 94158, USA

## Abstract

Complex I (CI) is the largest and most disease-associated component of the mitochondrial electron transport chain. While many diseases are linked to defects in specific CI subunits, the extent to which non-mitochondrial proteins contribute to CI function or disease is less clear. Here, we perform genome-wide CRISPR screens to identify regulators of CI abundance across its N, Q, and P modules, which mediate NADH oxidation, quinone reduction, and proton pumping, respectively. These screens identify THAP12 as a previously unrecognized transcriptional regulator of CI biogenesis. THAP12 loss selectively destabilizes CI and impairs oxidative ATP production. Mechanistically, THAP12 functions in the nucleus as a DNA-binding factor that directly activates genes required for CI assembly and iron-sulfur cluster maintenance, including NDUFAF3, NDUFAF4 and BOLA3. Patient-derived fibroblasts carrying THAP12 mutations exhibit conserved transcriptional defects and profound CI deficiency, establishing THAP12-associated neurodevelopmental disorder as a secondary mitochondrial CI disease. Finally, hypoxia rescues growth defects in THAP12-deficient cells, nominating low-oxygen therapy as a potential treatment strategy. Together, these findings identify THAP12 as a dedicated regulator of CI assembly and expand the genetic landscape of CI disease.

## INTRODUCTION

The mitochondrial electron transport chain (ETC) is an ancient system whose assembly and function require coordinated gene expression across the nuclear and mitochondrial genomes. Comprising four multi-subunit complexes (Complexes I-IV) in the inner mitochondrial membrane, the ETC oxidizes reduced cofactors (NADH, FADH_2_) and couples electron transfer to proton gradient generation for ATP synthesis. Beyond bioenergetics, ETC function supports essential biosynthetic reactions including pyrimidine synthesis, heme biosynthesis, and aspartate production. Among ETC components, Complex I (CI) is the largest and most implicated in disease^1^. It is assembled by over 20 dedicated factors and consists of 45 individual proteins that are organized into three functional modules: the N module (NADH oxidation), Q module (quinone reducing), and P module (proton pumping). A key consequence of this complexity is that loss of individual subunits or assembly factors typically destabilizes entire modules of the complex, making CI subunit abundance a sensitive proxy for complex integrity.

Mutations in CI subunits or assembly factors cause Leigh syndrome, a devastating pediatric neurological disorder characterized by bilateral symmetric brainstem lesions, progressive respiratory failure, and early death, for which no approved therapies exist^2–4^. We have shown that inhaled hypoxia extends lifespan nearly five-fold in the premiere mouse model of Leigh syndrome, the Ndufs4 knockout (KO) mouse. Late-stage hypoxia treatment can even reverse established disease^5,6^. Conversely, supplemental oxygen is catastrophic, causing death within hours to day^6,7^. This oxygen sensitivity reflects oxidative destabilization of the iron-sulfur containing N module and highlights CI module integrity as a mechanistically interpretable disease biomarker. These findings raise the question of what other conditions may cause secondary CI disease and whether they too are amenable to hypoxia therapy and vulnerable to hyperoxia.

While CI disease has been historically attributed to mutations in CI subunits and assembly factors, it is increasingly appreciated that genes without direct roles in CI can converge on CI dysfunction. For example, mutations in iron-sulfur cluster biogenesis genes (NFU1, BOLA3, FXN), the cardiolipin remodeling enzyme tafazzin (TAZ), and the mitochondrial kinase PINK1 all cause secondary CI loss through distinct mechanisms^8–14^. Together, these observations suggest that the scope of CI disease extends well beyond known disease genes, and motivated us to embark on a comprehensive search for factors that impinge on CI biology.

An intriguing feature of Complex I is that its abundance must be tuned to the energetic demands of the cell, yet it is unknown how gene expression of the structural subunits and assembly factors is regulated. While the PGC-1α/NRF1/TFAM pathways regulate broad mitochondrial biogenesis, no transcription factor has been shown to specifically control CI. The closest example is ZBTB11, a transcription factor that broadly regulates bioenergetic genes through its recruitment of NRF-2/GABP^15^. To address these gaps, we performed a FACS-based genome-wide CRISPR screen to uncover modifiers of CI, using subunit abundance as a readout of CI integrity. Screens assessing N, Q, and P module abundances were performed in parallel to distinguish general and module-level modifiers of CI. This approach identified THAP12 as a novel transcriptional regulator of CI assembly. These findings establish a new layer of CI regulation, expand the CI disease gene landscape, and identify THAP12 as a candidate disease target for hypoxia therapy.

## RESULTS

### FACS-based CI abundance screen recover known modifiers of CI biogenesis and assembly

To systematically identify genetic determinants of CI homeostasis, we developed an antibody-based flow cytometry assay designed to assess the abundance of the N, Q and P Modules using representative subunits (**Figure 1A**). To validate this approach, we subjected cells to chloramphenicol treatment (mitochondrial translation inhibitor) or hyperoxia exposure (50% O_2_), perturbations known to decrease CI abundance. Both conditions resulted in the expected depletion of CI modules when measured by flow cytometry, confirming the sensitivity of our detection method (**Figure 1B, 1C, and Figure S1A-D**). As a control for overall mitochondrial abundance, we also assessed the levels of citrate synthase (CS), a TCA enzyme. CS levels were not affected by chloramphenicol treatment or hyperoxia exposure. These results demonstrate that antibody-based flow cytometry provides a robust platform for the assessment of CI modifiers at scale.

**Figure 1.**
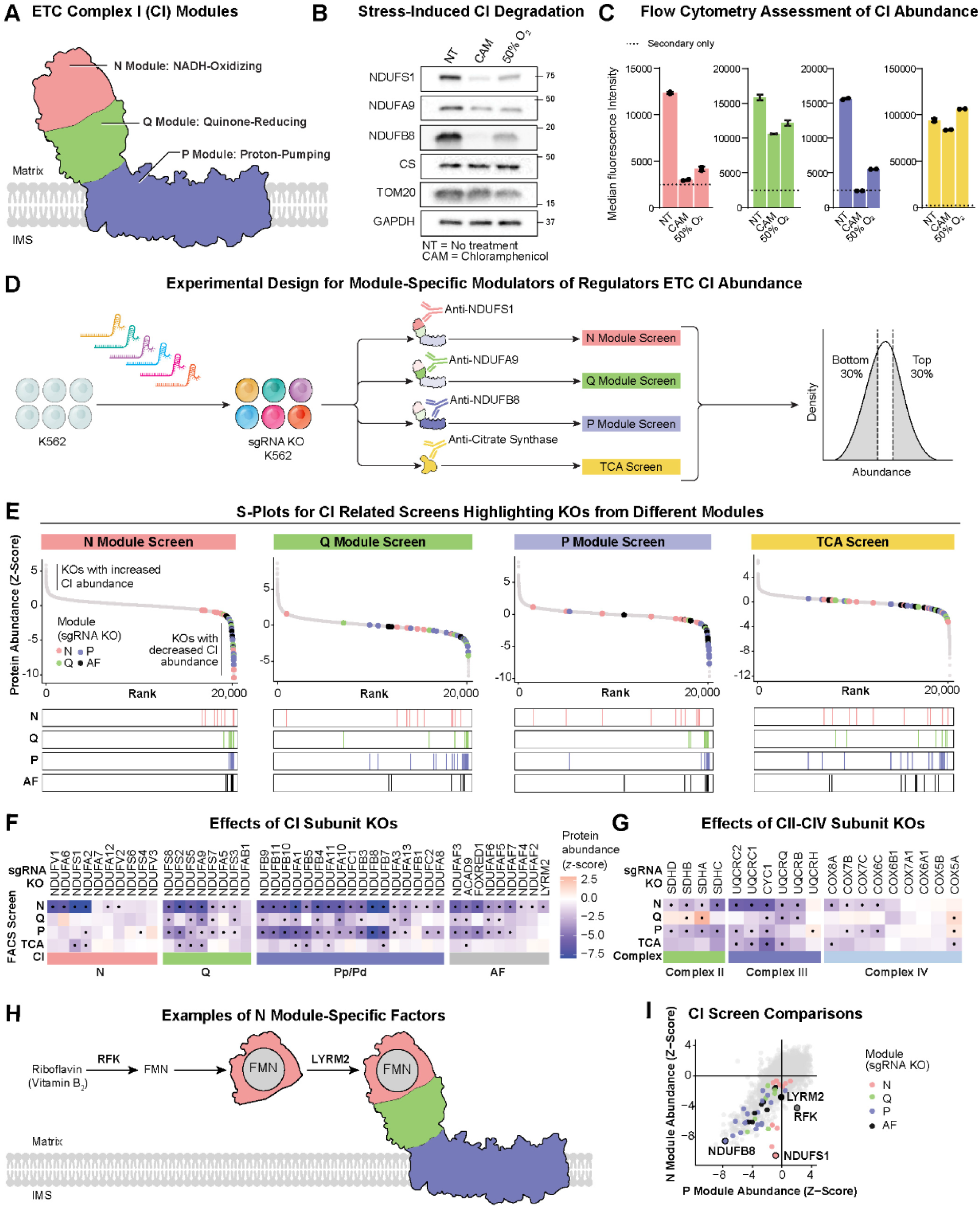
Modifiers of respiratory complex I stability revealed by FACS-based abundance screen. (A) Complex I (CI) structure is organized into modules based on partitioned functions. Electrons enter CI through the N module via NADH oxidation resulting in the reduction of quinone at the Q module. Free energy from these reactions supports proton pumping across the inner membrane at the P module. IMS = Inner membrane space. (B) SDS Western blot (WB) assessment of NDUFS1 (N module), NDUFA9 (Q module), NDUFB8 (P module) and CS (citrate synthase, TCA enzyme) abundance following mitochondrial translation inhibition with chloramphenicol (CAM), or hyperoxia (50% O2) exposure. Representative blot of two independently repeated experiments. (C) Flow cytometry assessment of NDUFS1 (N module, pink), NDUFA9 (Q module, green), NDUFB8 (P module, blue) and CS (yellow) protein abundance following mitochondrial translation inhibition or hyperoxia (50% O2) exposure. Bar represents mean +/- S.D. for a representative experiment shown (N = 2 technical replicates; one of two biological replicates shown). (D) Schematic depiction of genome-wide FACS-based CRISPR screening approach for uncovering genetic modifiers of CI abundance. A library containing 80,000 sgRNA guides (4x sgRNAs/gene) was transduced into WT K562 cells alongside Cas9. Transduced cells were selected, fixed, and immunostained for each CI module in parallel. Immunolabeled cells in the bottom or top 30% were sorted and sequenced to assess sgRNA enrichment. Sequencing results were analyzed using MAGeCK. (E) S-Rank plots depicting the effect size (*z*-score = (LFC_gene_ - LFC_mean.all.genes_)/S.D.) and rank for each gene across all performed screens. Data points for CI subunits are colored according to their respective structure module. (F) Heatmap showing the impact (z-score value) of individual CI subunits’ knockout or assembly factors’ (AF) knockout in the CI (N, Q, P) or CS screens. Genes scoring with a false discovery rate (FDR) < 0.1 are denoted with a dot. (G) Same as F but for KOs of CII, CIII, and CIV subunits. (H) Schematic depicting roles for RFK and LYRM2 in N module maturation and addition. (I) Impact of gene KOs on N module vs P module. Each dot is a single gene KO. Colored dots correspond to subunits of the N, P or Q module or to AFs. Knocking out AFs RFK and LYRM2 reduces N but not P module abundance.

Leveraging this approach, we performed genome-wide CRISPR knockout (KO) screens in K562 cells using the Brunello library to identify genetic modifiers of CI abundance^16^. We sorted populations based on the high and low protein abundance of NDUFS1 (N module), NDUFA9 (Q module), NDUFB8 (P module), or CS (TCA control), followed by deep sequencing to identify enriched sgRNAs associated with modified abundance for each protein (**Figure 1D, Figure S1E, Supplemental Table 1**). We identified many KOs that selectively impacted CI, but not CS. This included canonical CI assembly factors, such as ACAD9 or NDUFAF3 (**Figure 1E, 1F, and Figure S2A-D**). Nearly all CI subunit KOs led to the depletion of the N module, suggesting that N module stability is a sensitive indicator of overall complex integrity. In contrast, P module abundance was selectively diminished by KOs of P module subunits but remained largely unaffected by the loss of N module subunits. These module-specific depletion patterns are consistent with previously reported proteomics of individual CI subunit KO cells^17^. We also observed that CI abundance was sometimes affected by the loss of subunits from Complexes II, III, and IV (CII-CIV) with the depletion of CII and CIII subunits exerting the most pronounced impact (**Figure 1G**).

Beyond regulators of overall CI abundance, we also identified several module-specific factors such as *RFK* and *LYRM2* (**Figure 1H, 1I)**. Both factors are directly implicated in N module maturation: RFK (Riboflavin Kinase) is the essential kinase responsible for converting riboflavin into flavin mononucleotide (FMN), the critical redox cofactor embedded within the N module^18,19^. Once the N module is pre-assembled and flavinated, LYRM2 serves as a late-stage assembly factor responsible for the integration of the N module with the Q module in the holo-complex^20^. Collectively, these data demonstrate that our screening strategy not only identifies general regulators of CI abundance but also provides insights into the module-level biology of CI.

### FACS-based CI abundance screen uncover negative regulators of CI biogenesis

Importantly, our screens also identified gene KOs that increased the abundance of Complex I subunits (**Figure S3**). There was significant concordance with a previous screen performed in glucose vs galactose that identified KOs that increased cell growth in galactose conditions during which cells must rely exclusively on ETC-dependent oxidation of fuels (since glucose is not available for anaerobic glycolysis)^21^. Future studies will focus on testing gene KOs that increase CI abundance to determine whether they also affect CI activity and might serve as therapeutic targets to boost CI function for rare genetic mitochondrial disease or common conditions associated with CI defects such as Parkinson’s disease.

### Complex I abundance screens discover non-mitochondrial positive regulators of complex I

A key goal of our screens was to identify novel, non-mitochondrial modifiers of CI that cause human disease. To more specifically narrow in on such genes, we ran our screen results through a number of filtering steps (**Figure 2A-C**). First we removed any shared hits with the CS screen to eliminate general regulators of mitochondrial abundance. Next we prioritized genes that impacted CI but not other mitochondrial complexes. To this end, we compared hits from our N module screen to hits from a recent CRISPR screen for CIV modifiers^23^ (**Figure 2B**). This prioritization yielded 205 genes that specifically impacted NDUFS1 abundance without significantly affecting CS or CIV levels. Notably, 107 of these genes (52%) encode non-mitochondrial proteins, highlighting the extensive extra-mitochondrial influence over CI homeostasis (**Figure 2C**). Our prioritized list successfully recovered several known non-mitochondrial modifiers of CI. These included SLC52A2, a plasma membrane riboflavin transporter whose loss limits the FMN availability required for N module stability^24–26^; NCOA4, a selective autophagy receptor for ferritin (ferritinophagy) essential for mobilizing iron stores for iron-sulfur cluster biogenesis^27,28^; and CLUH, an RNA-binding protein known to coordinate the localization and expression of select mitochondrial-targeted mRNAs^29,30^. The identification of these diverse factors confirms that our screening strategy is very sensitive to the wide range of physiological inputs required to maintain CI integrity.

**Figure 2.**
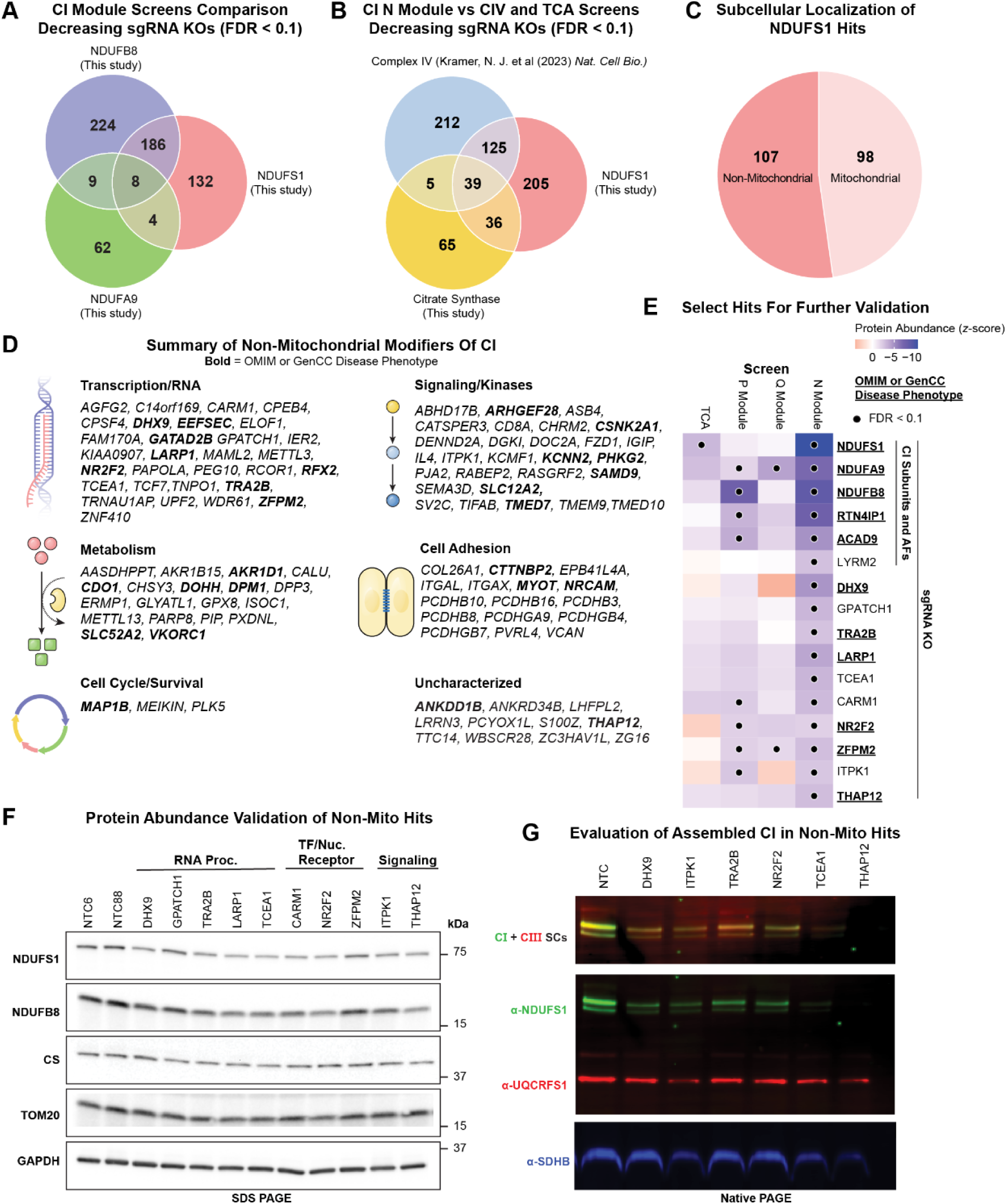
Complex I abundance screens discover non-mitochondrial positive regulators of complex. **I.** (A) Comparison of CI screen hits highlighting sgRNA KOs resulting in decreased module abundance. Hits also identified in the CS screen were removed before comparison (evaluated by an FDR < 0.1 derived from MAGeCK analysis). (B) Comparison of N module screen to CS screen and CIV abundance screen (Kramer, N.J. et al. (2023) *Nat. Cell Bio*). (C) Distribution of N module hits identified in (B) according to subcellular localization of their protein products. (D) Summary of non-mitochondrial hits identified in (C) grouped based on their expected function. Genes in **bold** are included in OMIM or GenCC, indicating a documented human disease phenotype. (E) Heatmap depicting screen results for select hits alongside known CI subunits and assembly factors for comparison. (F) SDS WB validation of CI screen hits prioritized in (E). (G) Blue Native PAGE WB validation of CI screen hits.

To prioritize clinically-relevant screen hits, we cross-referenced non-mitochondrial N module modifiers with the Online Mendelian Inheritance in Man (OMIM) and Gene Curation Coalition (GenCC) databases and found that 44 of the non-mitochondrial hit genes coincide with known human disease phenotypes (**Figure S4**). Roughly half of these are associated with neurodevelopmental disorders. For the majority of these conditions, there is no currently documented link to mitochondrial dysfunction or ETC deficiency. The identification of these genes in our screen suggests that a relatively large subset of neurodevelopmental pathologies may be driven by unrecognized defects in CI biogenesis or stability, offering a potential mechanistic explanation for previously idiopathic clinical presentations and warranting further functional exploration.

Building on these analyses, we selected 12 non-mitochondrial modifiers, most of which associated with human disease, for biochemical validation. This subset spanned diverse biological functions, including RNA processing enzymes, transcription factors, and cell signaling proteins (**Figure 2D, 2E**). We first confirmed the impact of these hits on the steady-state levels of CI subunits via SDS-PAGE and western blotting. In nearly all tested cases, KO of the candidate gene resulted in a reduction of CI subunit abundance, validating the results of our FACS-based screen (**Figure 2F**).

To determine if these reductions in subunit levels translated to defects in the assembly status of CI, we performed digitonin-based Blue Native PAGE (BN-PAGE) on a subset of six high-priority non-mitochondrial hits. This approach allowed us to visualize the integrity of both the fully assembled CI and its higher-order supercomplexes. Consistent with our subunit-level data, all tested KOs led to a marked decrease in the abundance of assembled CI and CI-containing supercomplexes. Among these candidates, THAP12 KO resulted in the most dramatic depletion of assembled CI (**Figure 2G**). Given this robust phenotype, the lack of previously documented roles for THAP12 in mitochondrial bioenergetics, and its link to neurodevelopmental disease, we prioritized this factor for subsequent mechanistic and disease-linkage characterization.

### THAP12 is required for maintaining Complex I integrity

THAP12 ranked within the top 1% of all protein-coding genes for knockouts resulting in CI depletion, marking it as a top non-mitochondrial regulator of CI abundance (**Figure S5A-D**). To determine whether its effect was specific to CI, we performed label-free whole-cell proteomics on THAP12 KO K562 cells. CI subunits were among the most significantly depleted proteins in response to THAP12 loss (**Figure 3A, Figure S6, Supplemental Table 2**). Notably, CI protein depletion extended beyond core structural subunits and included the CI assembly factors NDUFAF3 and NDUFAF4. In all, 30 of the 43 detected CI subunits, across all three modules, were significantly depleted (**Figure 3B**).

**Figure 3.**
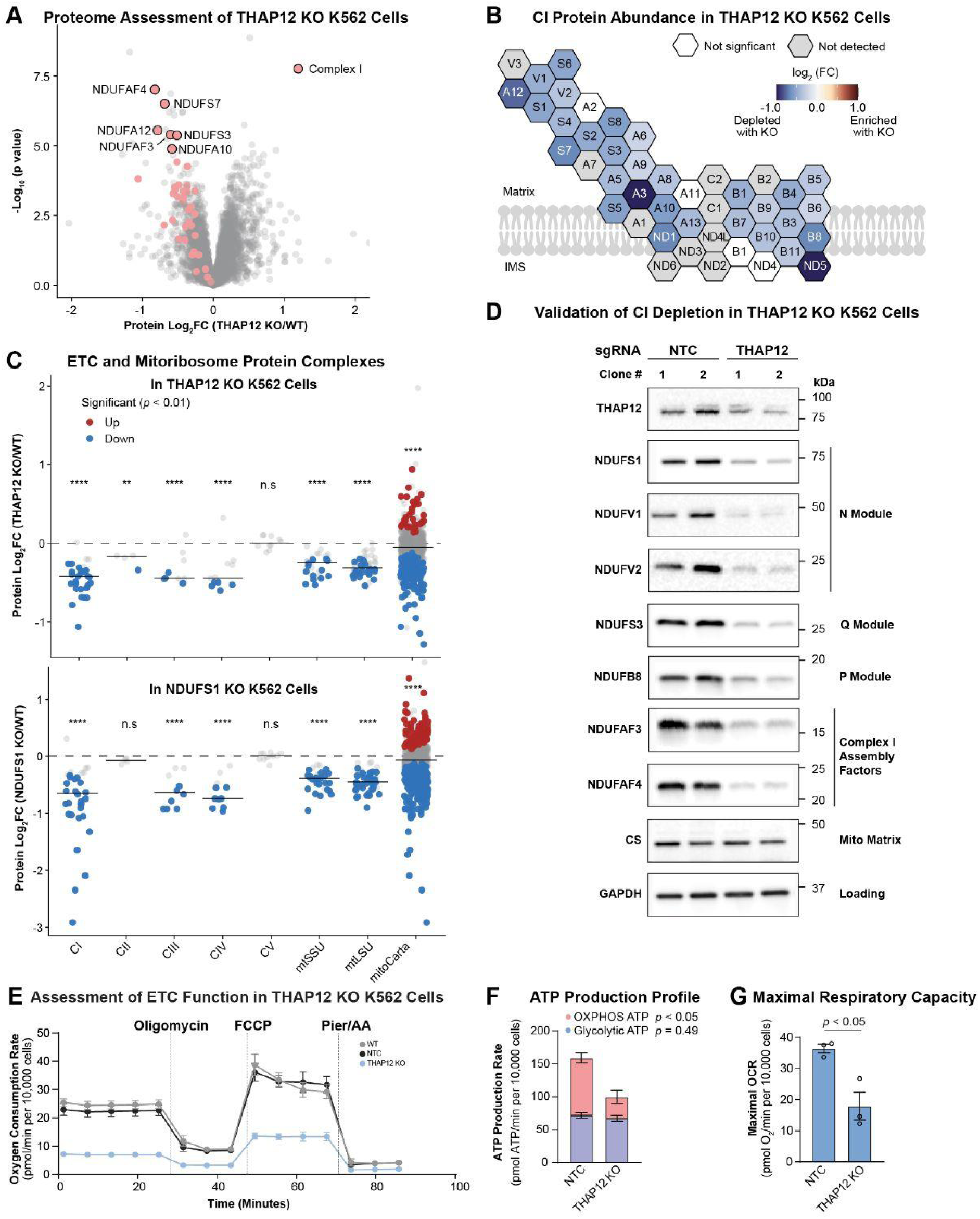
THAP12 is required for maintaining Complex I integrity. (A) Volcano plot highlighting CI subunits. Labels are included for the most significantly changed CI subunits. (B) Schematic depiction of CI overlaid with proteomics data from THAP12 KO K562 cells. (C) Jitter dot plot depicting abundance of individual ETC subunits and mitoribosome subunits grouped by their corresponding protein complex. Median log2FC for each complex is depicted with a horizontal line and evaluated against all changed proteins in the dataset and alongside all mitoproteins (annotated based on presence in MitoCarta 3.0). Comparisons were evaluated using wilcoxon test. Top: THAP12 KO vs NTC K562 cells (N = 3 biological replicates; Bottom: NDUFS1 KO vs NTC K562 cells (N = 2 biological replicates; Differences in median abundance of mitochondrial protein groups was compared to background proteins in the dataset using the Wilcoxon Rank Sum Test. Statistical significance is indicated as follows: * p < 0.05, ** p < 0.01, *** p < 0.001, and **** p < 0.0001. (D) Validation of CI subunit and assembly factor depletion in monoclonal THAP12 KO K562 cells. (E) Representative Seahorse respirometry trace from one of three experiments assessing oxygen consumption rate in THAP12 KO K562 cells. Points represent mean ± SD. (F) ATP production rate from oxidative phosphorylation and glycolysis, as derived from Seahorse respirometry. Bars represent mean ± SEM (N = 3 biological replicates). (G) Maximal respiratory capacity elucidated with FCCP. Bars represent mean ± SEM (N = 3 biological replicates). Unpaired t-tests were performed for evaluating differences between NTC and THAP12 KO in F and G.

CI was more severely affected than other ETC complexes and mitochondrial ribosomal proteins in THAP12 KO cells (**Figure 3C, top panel)**. To determine if the milder loss of these other complexes was secondary to the CI loss, we analyzed the proteomic profile of NDUFS1 KO cells that have a primary CI defect. The loss of this core CI subunit was sufficient to recapitulate the secondary depletion of CIII, CIV, and mitoribosome proteins (**Figure 3C, bottom panel**), suggesting that the milder defects in other ETC complexes in THAP12-deficient cells likely arise downstream of a primary failure in CI stability. Western blot analysis confirmed that THAP12 KO resulted in a dramatic loss of CI structural subunits, as well as assembly factors NDUFAF3 and NDUFAF4 **(Figure 3D**).

To determine if the observed loss of CI stability translated into functional defects, we assessed the bioenergetics of THAP12 KO K562 cells. Loss of THAP12 resulted in impairment of ETC-driven respiration (**Figure 3E**). This respiratory deficiency coincided with a reduced rate of mitochondrial ATP production leading to a significant decrease in the total cellular ATP production rate (**Figure 3F**). Furthermore, THAP12 KO cells displayed a decreased maximal respiratory capacity, a finding consistent with the near-total loss of CI subunits identified in our proteomic analyses (**Figure 3G**). Collectively, these physiological findings offer orthogonal functional validation for a significant CI impairment in THAP12-deficient cells.

### THAP12 regulation of Complex I biogenesis is independent of PKR signaling

THAP12, originally identified as PRKRIR or P52rIPK, was initially characterized as a regulator of the protein kinase R (PKR) branch of the Integrated Stress Response (ISR)^31,32^. In this model, THAP12 binds to and inhibits DNAJC3 (P58rIPK), an inhibitor of the stress kinase PKR. By sequestering DNAJC3, THAP12 de-represses PKR, leading to the phosphorylation of eIF2a and subsequent activation of the ISR (**Figure S7A**). Given the known interplay between mitochondrial stress and the ISR, we investigated whether the CI depletion observed in THAP12-deficient cells was a secondary consequence of disrupted PKR signaling.

To test this, we generated THAP12, DNAJC3, and PKR KO K562 cells and assessed their impacts on CI abundance. While THAP12 loss resulted in dramatic CI deficiency, KO of PKR or DNAJC3 did not affect CI levels (**Figure S7B-C**). These results indicate that THAP12’s role in maintaining CI biogenesis is distinct from a previously described role in the PKR-dependent ISR, suggesting a novel, unappreciated mechanism.

### Nuclear THAP12 expression is sufficient to restore complex I biogenesis

To determine the site of THAP12 action, we first characterized its subcellular localization by immunofluorescence. Endogenous THAP12 was detected in both the nucleus and the cytoplasm of HeLa cells, suggesting the potential for roles in both compartments (**Figure 4A**). To resolve which localization is functionally relevant for CI integrity, we engineered THAP12-3xFLAG variants designed for compartment-specific expression. Nuclear THAP12 was generated by fusing a C-terminal SV40 nuclear localization signal (NLS) to the protein (**Figure 4B)**. To achieve cytoplasmic localization, we neutralized a predicted endogenous NLS and appended a nuclear export signal (NES); this dual-modification was necessary as NLS neutralization alone did not fully exclude THAP12 from the nucleus (**Figure 4C and Figure S8**).

**Figure 4.**
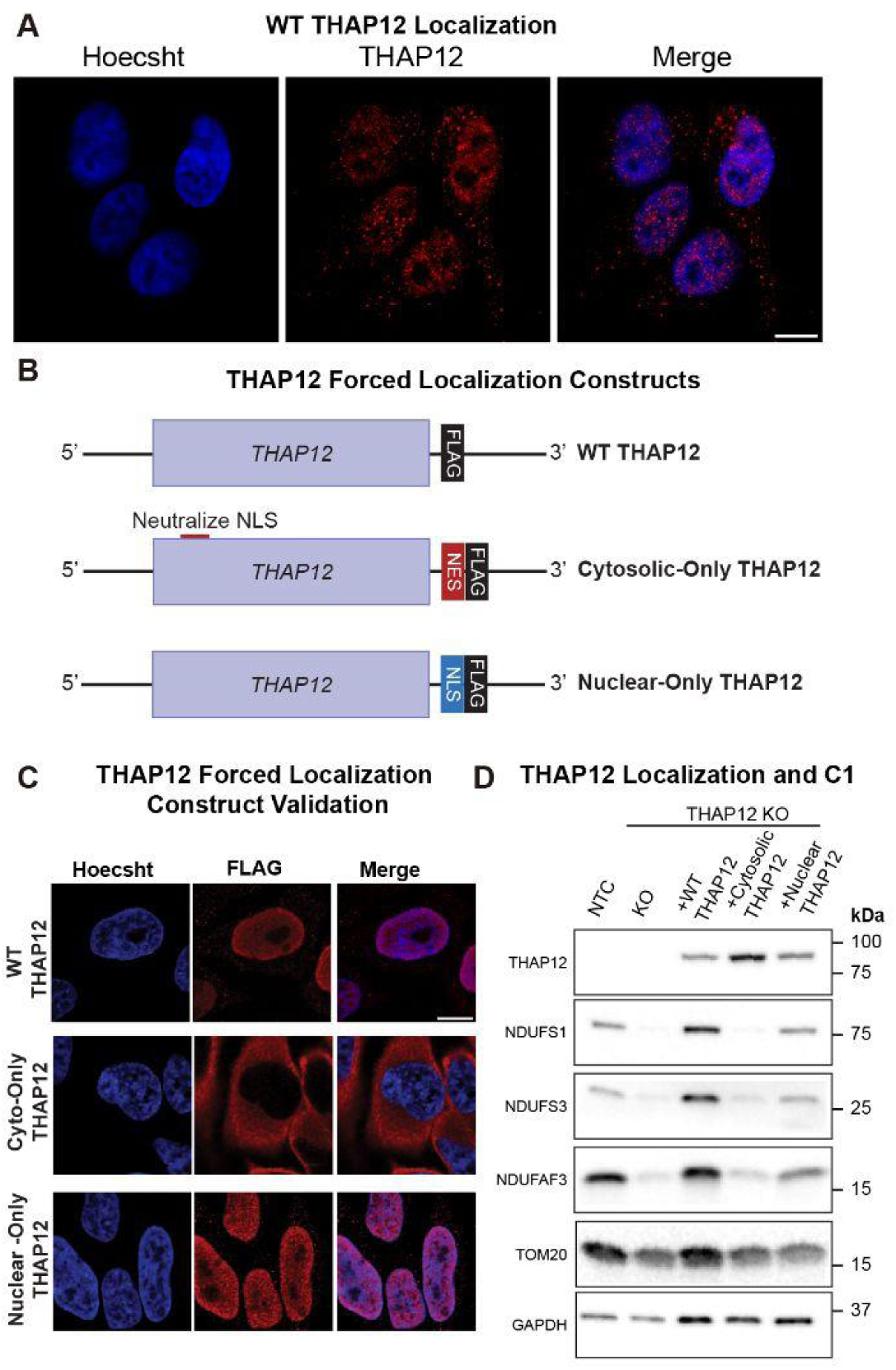
Nuclear THAP12 expression is sufficient to restore Complex I biogenesis. (A) Representative immunofluorescence confocal micrograph of WT HeLA cells depicting THAP12 subcellular localization. (B) Cartoon schematic of engineered THAP12 overexpression constructs for assessing nuclear and cytosolic functions. (C) Representative micrographs from engineered THAP12 OE HeLa cells validating correct forced localization. (D) SDS WB assessment of CI subunit abundance with forced localization of THAP12.

Upon stable re-expression of these variants in THAP12 KO cells, we observed that both wild-type (WT) and nuclear-targeted THAP12 (THAP12-NLS) successfully restored CI subunit abundance to near-WT levels (**Figure 4D**). On the other hand, the cytoplasmic-restricted variant failed to rescue the CI phenotype, indicating that the nuclear pool of THAP12 is the primary driver of CI homeostasis. These results demonstrate that THAP12 functions as a nuclear factor essential for the biogenesis of CI.

### THAP12 regulates expression of factors necessary for CI biogenesis through direct binding of their promoters

Given the requirement for the THAP12 DNA-binding domain in rescuing CI levels, we performed RNA-seq to identify the specific role for THAP12 in regulating gene expression. Differential expression analysis revealed a marked downregulation of several transcripts essential for CI biogenesis including NDUFAF3 and BOLA3 (**Figure 5A and Supplemental Table 3**).

**Figure 5.**
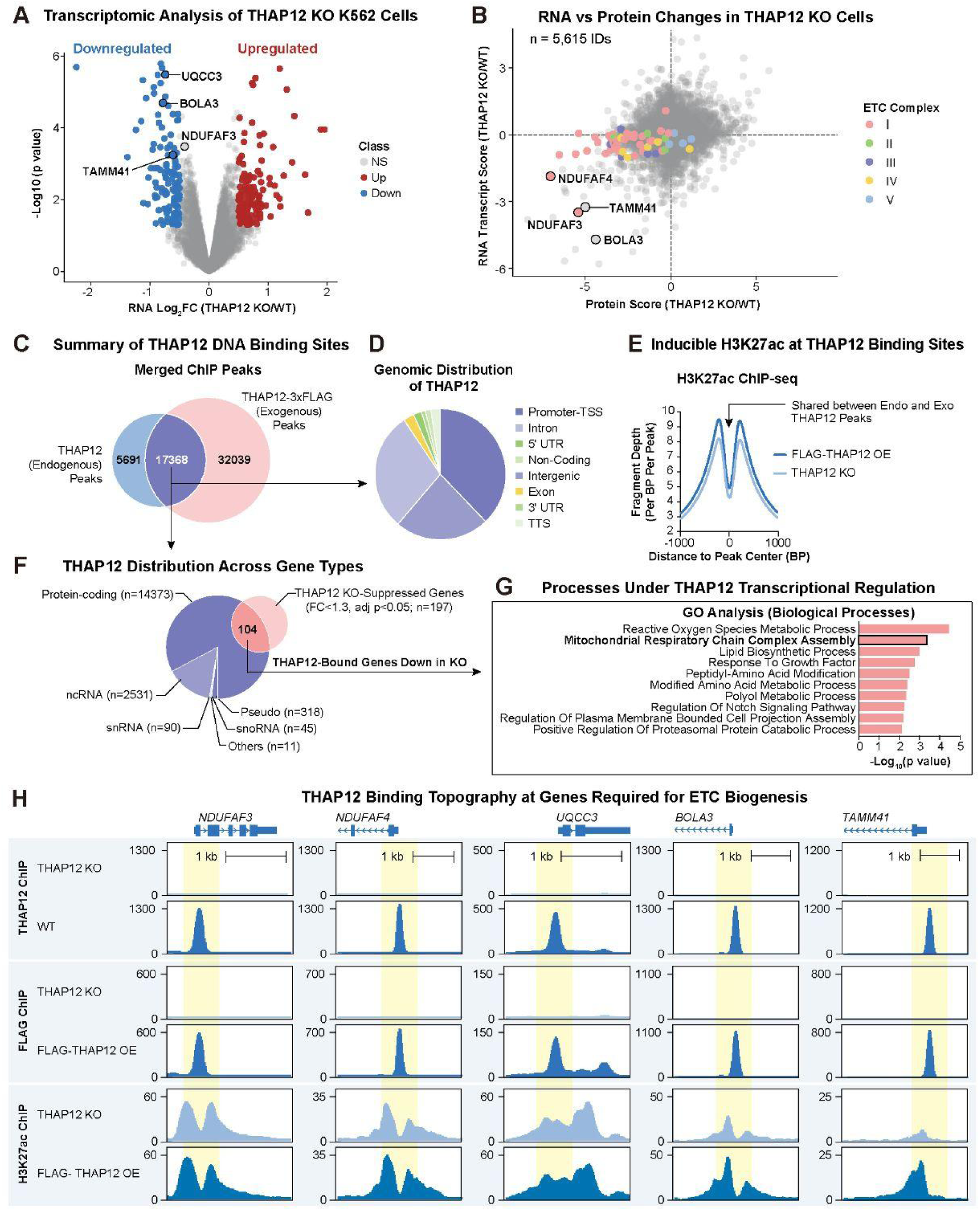
Nuclear THAP12 binds at the promoter of genes required for Complex I biogenesis. (A) Differential gene expression from RNAseq depicted as log_2_FC. (B) Comparison of differential gene expression score versus differential protein abundance score in THAP12 KO K562 cells. Scores were calculated as sign(logFC) × –log10(P value). Labeled dots are factors required for CI assembly. (C) Summary of THAP12 binding sites identified with ChIPseq from an endogenous THAP12 IP and THAP12-3X FLAG IP. (D) Characterization of THAP12 binding sites identified with ChIPseq. (E) Assessment of H3K27ac at THAP12 binding sites via H3K27ac ChIPseq. (F) Summary of THAP12 binding sites based on gene type. Protein-coding genes are further compared to suppressed genes identified by RNAseq in (A). (G) Summary of processes under THAP12 regulation, as identified by THAP12 binding with ChIPseq and suppressed gene expression with RNAseq. (H) THAP12 binding topography at genetic loci important for ETC biogenesis.

To identify high-confidence direct targets, we correlated the RNASeq and proteomics datasets from THAP12 KO vs WT cells. This revealed a striking concordance between mRNA and protein depletion for a subset of critical regulators of CI. Specifically, both the transcript and protein levels of BOLA3 and the NDUFAF3 were among the most severely depleted in the entire analysis, with NDUFAF4 also trending in the same direction (**Figure 5B**). This suggested that THAP12 is required for their transcriptional maintenance, providing a potential mechanistic link between nuclear THAP12 activity and CI integrity.

To identify the direct genomic targets of THAP12, we performed ChIP-seq in both wild-type and THAP12-3xFLAG expressing cells using anti-THAP12 or anti-FLAG. We focused the subsequent analysis on the overlapping set of peaks for these two antibodies (**Figure 5C**). Global analysis of THAP12 binding topography revealed that the vast majority of binding peaks were concentrated at the promoters of protein-coding genes (**Figure 5D**). By integrating parallel H3K27ac ChIP-seq data, we confirmed that these THAP12-bound loci are characterized by high levels of histone acetylation adjacent to the transcription start sites (TSS), a hallmark of transcriptionally active promoter regions (**Figure 5E**).

To refine this list into high-confidence regulatory targets, we prioritized genes that were both physically bound by THAP12 and differentially expressed in our transcriptomic dataset **(Figure 5F)**. This intersectional analysis identified a core set of 104 THAP12-regulated genes. Notably, Gene Ontology (GO) analysis revealed that “mitochondrial respiratory chain complex assembly” was one of the most significantly enriched biological processes within this group (**Figure 5G**). Among these direct targets were several essential CI biogenesis factors identified in our initial FACS screen, including NDUFAF3, NDUFAF4, BOLA3, and the cardiolipin biosynthesis factor TAMM41 (**Figure 5H and Figure S9**). Notably, mutations within these genes are each strongly linked to mitochondrial disease in patients. These results establish that THAP12 is a direct transcriptional activator of a specialized nuclear program required for the assembly and maintenance of CI.

### THAP12 disease mutations exhibit Complex I deficiency that is rescued by hypoxia

To determine the clinical relevance of our findings, we examined fibroblasts derived from patients carrying mutations in THAP12. THAP12 has recently been identified as a candidate gene for a neurodevelopmental disorder, inherited in an autosomal recessive manner (**Figure 6A**). The affected individuals described to date carry compound heterozygous variants, including a frameshift allele and a missense mutation. Both patients present with microcephaly, drug-resistant seizures, and significant neurodevelopmental delay^33^. Currently, both patients are diagnosed with Lennox-Gastaut syndrome. However, the molecular mechanisms driving their disease remains unclear. To assess if CI deficiency is a feature relevant to THAP12 disease, we performed a WB analysis of CI subunit abundance in patient fibroblasts from THAP12 patients. We observed dramatic depletion of CI subunits, as well as the assembly factor NDUFAF3 (**Figure 6B**) compared to their carrier parents. Furthermore, we observed a profound depletion of BOLA3 protein levels, mirroring the defect seen in our engineered KO cells.

**Figure 6.**
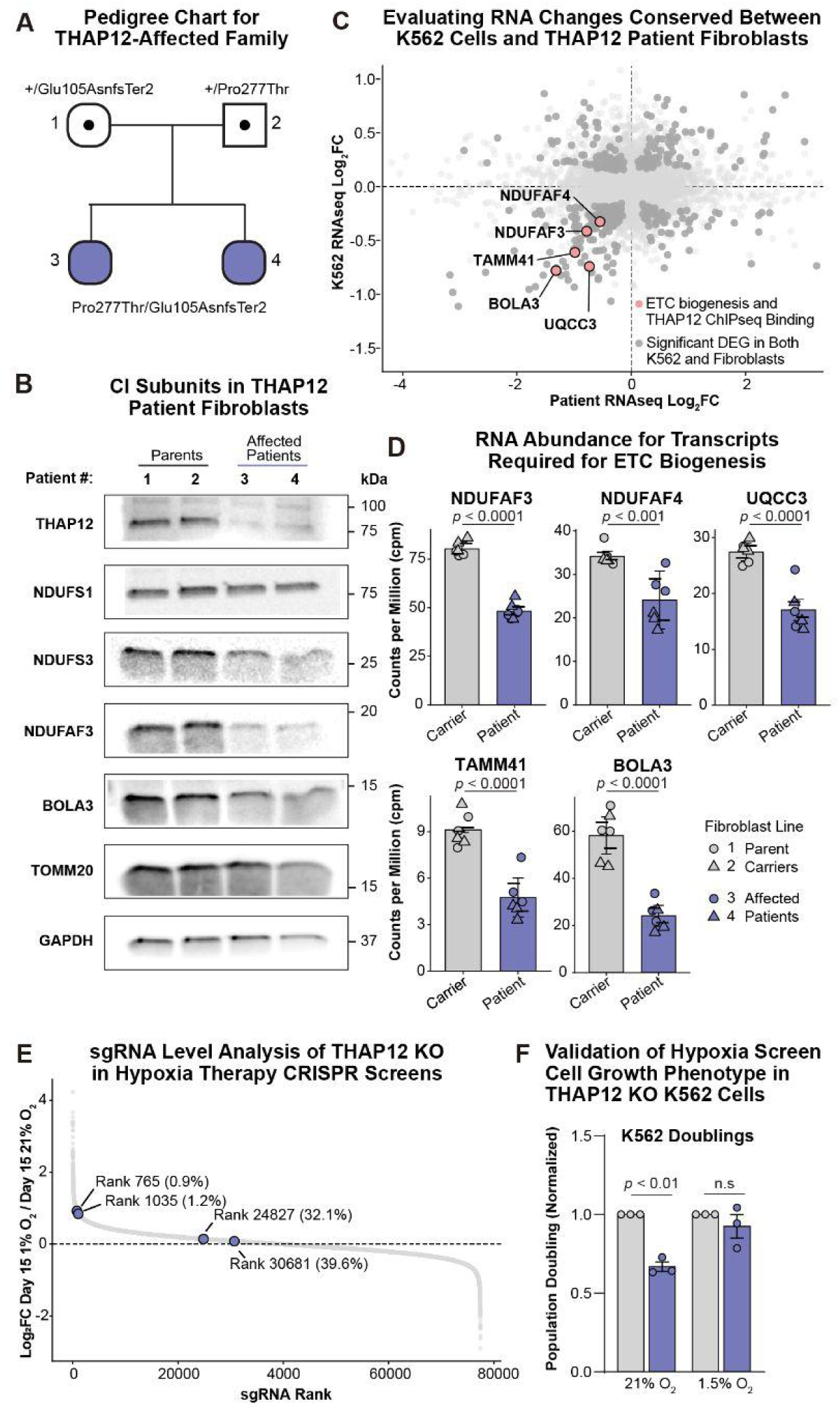
Complex I deficiency in THAP12 patient fibroblasts. (A) Pedigree chart depicting autosomal recessive inheritance pattern of THAP12 disease. (B) SDS WB assessment of CI subunits, assembly factor, and BOLA3 protein abundance. (C) Comparison of differential gene expression from THAP12 KO K562 cells vs. THAP12 patient fibroblasts. Differential gene expression is represented as log_2_FC. Labeled dots are factors required for CI assembly with observed THAP12 binding. (D) Normalized RNA abundance from labeled transcripts in (C). (E) sgRNA-level assessment of THAP12 KO in the hypoxia therapy screen (Jain 2020). (F) Cell growth assessment of monoclonal THAP12 KO cells cultured in normoxia (21% O_2_) or hypoxia (1.5% O_2_) for 3 days. Bars represent mean ± SEM (N = 3 biological replicates). Unpaired t-tests were performed for evaluating differences between NTC and THAP12 KO in F.

We next sought to confirm if these protein-level changes to CI were due to transcriptional impairment in the patient cells. We performed RNA-seq analysis on patient-derived fibroblasts and compared the results to our THAP12 KO K562 RNA-seq data (**Figure 6C and Supplemental Table 3**). This comparison revealed a highly conserved transcriptional signature; among the most significantly downregulated genes in both cell types were the core CI assembly and biogenesis factors NDUFAF3, NDUFAF4 and BOLA3, and TAMM41 (**Figure 6C, 6D**).Collectively, these data establish that the loss of THAP12-mediated transcriptional control of CI biogenesis is a conserved feature of the human disease, identifying THAP12-related disorder as a novel form of CI deficiency.

Given the severe CI deficiency observed in THAP12-deficient cells, we investigated whether this phenotype could be mitigated by hypoxia, a condition we have shown to rescue survival in several models of CI deficiency. We first performed a cross-comparison between our CI abundance screen and our previously generated “hypoxia therapy” screen designed in K562 cells to identify genes whose KO-induced growth defects are suppressed by low oxygen^34^. This analysis identified 88 shared hits that both destabilize CI and show a positive growth response to hypoxia (**Figure S10A-B**). Notably, this list included the THAP12 transcriptional targets NDUFAF3 and BOLA3. While THAP12 itself did not initially meet the stringent threshold for a hit in the hypoxia screen, a guide-level analysis revealed that it was strongly trending toward this category. Two of the four sgRNAs evaluated were clear hits, while a third guide showed no baseline growth defect in normoxia, precluding the measurement of a rescue effect (**Figure 6E, Figure S10C)**. To independently test whether THAP12-deficient cells benefit from low-oxygen environments, we compared cell proliferation of WT and THAP12 KO K562 cells under normoxic and hypoxic (1.5% O2) conditions. Due to a lack of an observable cell proliferation defect in the patient fibroblasts, we utilized K562 cells for this assessment. In normoxia, THAP12 KO cells exhibited a significant growth defect, consistent with their profound CI deficiency and reduced total ATP production (**Figure 6F, Figure S10D**). However, when cultured in hypoxia, this growth disparity was abolished, and the proliferation rate of THAP12 KO cells was no longer significantly slower than that of WT cells. This result recapitulates the “hypoxia therapy” response characteristic of other K562 models of CI deficiency. Collectively, these findings establish THAP12 as a novel regulator of ETC CI and identify hypoxia as a potential therapeutic strategy for THAP12-related secondary CI disease.

## DISCUSSION

In this work we use FACS-based genome-wide CRISPR screens to map the genetic regulators of Complex I (CI) homeostasis. By moving beyond traditional survival-based assays and utilizing a module-specific readout, we nominate new disease genes and therapeutic targets for CI disease. In doing so, we identified THAP12 as a transcriptional orchestrator of ETC Complex I.

Transcription factors (TFs) represent one of, if not the most underrepresented protein class related to mitochondria disease^2,35,36^ Outside of the classical mitochondrial TF TFAM and the core nuclear TF regulators NRF1 and GABPA, only ZBTB11 has been generally linked to ETC dysfunction and human disease^15^. Only one additional TF, E4F1, has been linked to secondary mitochondrial disease through dysregulated transcription of pyruvate dehydrogenase complex components.^37^. THAP12 is distinct from these broader regulators given its seemingly specialized regulation of CI assembly. Specifically, THAP12 drives the expression of NDUFAF3 and NDUFAF4, which coordinately manage matrix arm maturation integration into the pre-assembled P-module^38–40^. Furthermore, THAP12 directly controls BOLA3 expression, which delivers the [4Fe-4S] clusters essential for maintaining the structural and catalytic integrity of the matrix arm^9^. Lastly, loss-of-function mutations in any single one of these THAP12 target genes is independently sufficient to cause CI deficiency in our screens and independently linked to clinical mitochondrial disease. These observations underscore the critical input THAP12 has on CI integrity.

While we have established that the nuclear fraction of THAP12 is responsible for the transcriptional orchestration of CI assembly, its dual distribution across both the nucleus and cytoplasm suggests it may function as a dynamic regulatory node worthy of future investigation. Mechanistically, the subcellular trafficking of THAP12 is at least partially dictated by an internal bipartite nuclear localization signal (NLS), raising the intriguing possibility that an active regulatory mechanism exists to shift its localization to dynamically tune CI assembly status in response to metabolic or environmental demand. This regulation is reminiscent of what has been observed with longevity-associated FOXO TFs whose localization is regulated by Akt-dependent phosphorylation^41^. Because previous models have proposed that THAP12 interacts with DNAJC3 in a stress-dependent manner, a compelling avenue for future study will be to explore whether this interaction serves as the molecular switch that governs the nucleo-cytoplasmic partitioning of THAP12.

Beyond THAP12, our screens significantly broaden the conceptual scope of secondary mitochondrial disorders. Approximately half of the high-confidence hits from our N module screen encode proteins localized to mitochondria, many of which are linked to primary and secondary mitochondrial disease. The remaining half are non-mitochondrial proteins, and roughly half of these non-mitochondrial hits are associated with human disease phenotypes with little to no previous evidence of CI or ETC deficiency. The single largest disease category among these newly identified modifiers is neurodevelopmental disorders (NDDs). Although NDDs are widely heterogeneous and associated with a vast array of different genes, a growing body of literature over the past decade has consistently described underlying mitochondrial dysfunction in NDD patient cohorts^42–47^ Our findings provide additional experimental support for the hypothesis that a subset of these non-mitochondrial NDD genes may actually drive an unappreciated form of secondary CI disease.

Expanding our understanding of secondary mitochondrial disease driven by CI deficiency allows us to broaden the clinical scope of existing biomarkers and targeted therapeutics. Well-established metabolic (e.g. lactate, B-HB) or imaging (e.g. T2-weighted MRI hyperintense lesions) biomarkers used to monitor primary CI diseases could be readily adopted to track disease progression and evaluate therapeutic efficacy in patients with secondary disorders. Similarly, interventions pioneered to treat primary CI defects including rapamycin, and hypoxia therapy, could be repurposed for these conditions.^5,6,48^ For example, our work nominates THAP12 deficiency as a secondary CI disease that may be amenable to rescue by hypoxia or our newly developed HypoxyStat drug^49^. Of note, such secondary causes of CI disease may also have additional non-mitochondrial pathologies and therefore may display intermediate rescue effect sizes. Nonetheless, it expands the potential applicability of such therapies to hundreds of more disease genes beyond direct genetic lesions in CI subunits or assembly factors.

Additionally, the opposite side of our CRISPR screens identifies gene KOs that increase CI stability and may thus serve as therapeutic targets for treating CI disease. A prominent example includes *FASTKD1*, whose loss triggers a robust increase in both N and P module abundance. FASTKD1 was recently identified by others as a negative regulator of CI abundance through regulation of the mitochondrially-encoded CI transcript, *ND3*^50,51^. This example represents one of many CI negative regulators identified in our work. Future work should explore therapeutic potential for targeting these hits for treating diseases associated with CI deficiency.

In summary, our genome-wide screen mapped a comprehensive compendium of both enriching and destabilizing genetic modifiers of CI abundance with distinct module-level resolution. We successfully leverage the utility of our dataset to assign a crucial functional role to the previously uncharacterized protein THAP12. Based on this work, we propose that THAP12 acts as a specialized nuclear transcription factor that orchestrates the biogenesis of CI through the targeted regulation of key downstream pathways governing inner mitochondrial membrane maintenance, mitochondrial iron-sulfur cluster delivery, and direct CI assembly. Crucially, by demonstrating a robust CI deficiency at both the transcript and protein levels in primary fibroblasts from patients carrying mutations in this ultra-rare disease gene, we have validated the predictive power of our FACS-based screening strategy to uncover cryptic and challenging clinical pathologies. Future work should focus on defining the broader physiological roles of THAP12 alongside the precise regulatory nodes that control its expression and function. Ultimately, this study opens new avenues for studying, diagnosing, and treating primary and secondary CI disease.

## METHODS

### Cell Culture

K562 cells (ATCC, CCL-243) were grown in DMEM (11995, Thermo Fisher) containing high glucose (25mM) and pyruvate (1.25 mM). DMEM was also supplemented with Penicillin/Streptomycin (Thermo Fisher, 15140122) and 10% FBS (Corning, 3580). Cells used for CRISPR-Cas9 screening were also supplemented with 50 µg/mL uridine (Sigma Aldrich, U3003) to preserve cells with dysfunctional mitochondria.

Human dermal fibroblasts (Coriell; Patients: GM27990 and GM27993; Parent: GM27995 and GM27997) were cultured in EMEM (Sigma-Aldrich, M4655; with Earle’s salts and L-glutamine) supplemented with non-essential amino acids (Sigma-Aldrich, M7145), 1% Penicillin/Streptomycin, and 15% FBS.

Cells were cultivated in a humidified incubator maintained at 37 °C with 5% CO2. Hyperoxia and hypoxia experiments were performed using a similar incubator infused with 50% O_2_ or 1.5% O_2_ CO_2_.

### Genome-wide CRISPR Cas9 Screens

K562 cells (ATCC) were maintained in magnetic spinner flasks stirred at approximately 110 RPM throughout the duration of the screen. Cells were transduced with human lentivirus containing both Cas9 and the genome-wide Brunello library at a target MOI of 0.2 (Addgene). 400e6 cells were resuspended with supplemented DMEM (11995) containing 4ug/mL polybrene (TR-1003, Sigma Aldrich) at a concentration of 1.5e6/mL. Cells with polybrene were distributed in 24-well plates at a volume of 2mL/well before adding 45uL lentivirus/well. Cell plates containing lentivirus were centrifuged at 1,000xG for 2h at 32 °C. Approximately 18h following infection, infection media was replaced with DMEM containing 2ug/mL puromycin and cells were transferred to spinner flasks for the selection stage. A small portion of transduced cells were maintained in a 6-well plate and counted daily during selection to assess transduction efficiency.

Puromycin selection was carried out for 5 days. Cells were passaged once during selection when cells were nearly confluent. At 5 days post-infection, 80e6 cells were seeded in each flask according to their respective screen condition. Screening flasks were maintained in 21% O2 with the exception of the hyperoxia screen which was maintained at 50% O2 for the duration of the screen.

At screen day 3, 80e6 cells were passaged from each screening flask and further maintained until day 6. On day 6, samples were fixed with methanol as described in ref. (https://doi.org/10.1016/j.celrep.2022.111508). Briefly, 150e6 cells were pelleted and washed with ice-cold PBS. After washing, pelleted cells were fixed by slowly adding ice-cold methanol (1mL/10e6 cells) while gently vortexing. Cells were left on a rotator at -20 °C for 30 minutes. Samples were then immediately stored at -80 °C until sorting.

To prepare for intracellular antibody staining, fixed cells were rehydrated by adding 10 mL cold PBS to 10 mL of cells. Cells were pelleted at 300xG and resuspended with FACS blocking buffer (0.4% Triton X-100, 5% Normal Donkey Serum, 0.5% BSA) and incubated at 4 °C for 1h. After blocking, blocking buffer was replaced with primary antibody and incubated at 4 °C overnight.

Following overnight primary antibody staining, samples were washed three times with 0.4% PBST wash buffer. After the third wash, wash buffer was replaced with secondary antibody solution. Samples were incubated at room temperature on a rotator in the dark for 1.5h. Following secondary incubation, samples were washed three times with wash buffer. Finally, samples were resuspended with PBS containing 1 mM EDTA before filtering and sorting.

Cells were sorted at 10,000 events/s on a FACS Aria System fitted with a 70 um nozzle. Cells were gated based on fluorescence and approximately 200 cells per sgRNA were collected in each bin. Collected cells were pelleted and lysed for genomic DNA extraction (Machery Nagel). Sequencing libraries were prepared

### SDS PAGE

Protein lysates used for immunoblotting were prepared in RIPA lysis and extraction buffer (Thermo Fisher, PI89901) supplemented with a cocktail of protease inhibitors (Roche 11697498001). Lysates were clarified via centrifugation at 14,000xG for 15m. The resulting supernatant was transferred to a new tube, quantified, and abundance-normalized using bicinchonic acid (BCA) assays (Thermo Fisher, A55860). Clarified lysate was mixed with 6X Laemmli buffer and boiled at 95C for 5min before SDS-PAGE.

SDS-PAGE was performed using Mini-PROTEAN TGX pre-cast gels. All samples were run at a constant voltage of 200V, unless otherwise noted. Proteins were then transferred onto PVDF membranes using the semi-dry Trans-blot Turbo (BioRad, 1704157) system. Membranes were blocked with 3% non-fat milk in TBST buffer for 1hr at room temperature while rocking. After 1h, blocking buffer was replaced with blocking buffer containing primary antibody. Membranes were incubated with primary antibodies overnight at 4C. The following day, membranes were washed three times with TBST before incubating with blocking buffer containing the corresponding secondary antibody. Membranes were incubated with secondary antibodies for 1h at RT. Membranes were washed three times to remove unbound secondary antibody before addition of peroxidase substrate (Thermo Scientific, 32106) for chemiluminescent visualization and imaging. Chemiluminescence was imaged with using the Azure 600 Western blot imaging system (Azure, AZI600-01). All experiments were conducted with at least two biological replicates. One representative western blot is shown for each experiment.

### Blue Native PAGE

Fifty million K562 cells were resuspended in mitochondrial isolation buffer (0.32 M sucrose, 1 mM EDTA, 10 mM Tris–HCl, pH 7.4) with 1×cOmplete protease inhibitor (Roche) on ice and homogenized with ten strokes of a glass dounce homogenizer. Lysates were centrifuged at 1,000g to pellet unprocessed cells and nuclei, and the supernatant was re-centrifuged at 13,000g to enrich mitochondria. Pellets were stored at –80 °C until BN-PAGE processing. BN-PAGE was adapted from published protocols53.Mitochondria were resuspended at 10 μg per μg in native PAGE sample buffer and solubilized with 4 μg digitonin (Sigma) per microgram mitochondria for 10 min on ice, followed by centrifugation at 16,000g for 30 min at 4 °C. Supernatant was mixed with Coomassie Blue G loading buffer (1:3 volume) and supercomplexes resolved on native PAGE 3–12% Bis–Tris gradient gels (Thermo Fisher Scientific) using dark blue cathode buffer (150 V, 30 min), then light blue cathode buffer (300 V, 90 min). Proteins were transferred to methanol-activated PVDF membranes at 100 V for 60 min and blocked with 5% BSA in TBST before immunoblotting.

### Immunoblotting

After blocking, antibodies are prepared in 5% non-fat milk (SDS WB) or 5% BSA (BNGE WB). All primary antibodies were diluted 1:1000 unless otherwise notes. The primary antibodies used were NDUFS1 (Abcam, ab169540), NDUFA9 (Abcam, ab14713), NDUFB8 (Abcam, ab192878), Citrate Synthase (Cell Signaling Technology, 14309), SDHB (Abcam, ab178423), TOM20 (Cell Signaling Technology, 42406), GAPDH (Cell Signaling Technology, 2118S), UQCRFS1 (Abcam, ab14746), PRKRIR/THAP12 (Bethyl Laboratories, A300-586), NDUFV1 (Proteintech, 11238-1-AP), NDUFV2 (Abcam, ab183715), NDUFS3 (Genetex, GTX105835), NDUFAF3 (Thermo Scientific, PA5-57250), NDUFAF4 (Abcam, ab191414), BOLA3 (1:500 dilution, Sigma-Aldrich, HPA046393), PKR (Cell Signaling Technology, 12297S), and p58IPK/DNAJC3 (C56E7) (Cell Signaling Technology, 2940S). Membranes were incubated with primary antibodies and gentle agitation at 4 °C. Membranes were washed 3 times with TBST for 5 min each wash. Washed membranes were incubated with horseradish peroxidase-linked (Cytiva, NA934) or fluorescent secondary antibodies (LICRO, 92632211) for 1h at room temperature with gentle agitation. Membranes were washed 3 more times before ECL (Thermo, 32209) exposure and imaging.

### Whole cell proteomics

Cell pellets (approximately 10-12 million K562 cells) were transferred to ice and immediately resuspended in 800 µL of proteomics solubilization buffer (5% SDS, 50 mM TEAB, 1X protease inhibitor), then vortexed until fully homogenized. Samples were incubated on a rotator in a 4°C cold room for 30 min, with an additional vortexing step at 15 min to ensure complete mixing. Lysates were then sonicated at high power for ten cycles of 30 s on/30 s off. Insoluble material was removed by centrifugation (14,000 × g, 15 min, 4°C), and the resulting supernatant was transferred to a new tube. Protein concentration was estimated by BCA assay on a 1:20 dilution of each lysate.

Protein digestion was performed using a Protifi suspension trapping (S-Trap) workflow. Disulfide bonds were reduced with reductant (55°C, 15 min) and free thiols alkylated with alkylating reagent (room temperature, 10 min). Samples were acidified to quench the reaction and combined with binding/wash buffer containing methanol. Samples were then loaded onto S-Trap columns in 600-µL aliquots and trapped by centrifugation (10,000 × g, 30 s), repeating until all material had passed through the matrix. Columns were washed four times with 400 µL binding/wash buffer (10,000 × g, 30 s per wash). Trapped proteins were digested with trypsin at a 1:10 (protease:protein, w/w) ratio diluted in the digestion buffer for 2 h at 47°C.

Peptides were eluted sequentially with digestion buffer, 0.2% formic acid in water, and 50% acetonitrile in water (80 µL each, with centrifugation at 10,000 × g for 1 min after each addition). Pooled eluates were dried by vacuum centrifugation and reconstituted in 5% acetonitrile/0.1% formic acid prior to LC-MS analysis.

### Immunofluorescence Confocal Microscopy

HeLa cells (50,000 cells) were seeded onto 12 mm glass coverslips in a 24-well plate and cultured overnight. Cells were washed once with PBS and fixed with 4% paraformaldehyde for 10–15 min at room temperature, followed by three 3 min washes with PBS. Fixed cells were blocked and permeabilized in a blocking and permeabilization solution (3% BSA, 1% glycine, 10% normal donkey serum, and 0.1% Triton X-100 in PBS) for 30 min at room temperature.

Primary antibodies were diluted in blocking solution and applied to cells at the following concentrations: rabbit anti-FLAG (1:500) or rabbit anti-PRKRIR/THAP12 (1:500). Coverslips were incubated with primary antibodies overnight at 4 °C. Cells were washed three times for 3 min each with PBS containing 0.1% Triton X-100 (PBST). Secondary anti-rabbit Cy3 (1:1000) antibody was diluted in blocking solution and applied to cells for a 1h incubation at room temperature. After secondary incubation, coverslips were incubated with Hoechst (1 µg/mL) diluted in PBST for 10m to label nuclei. Coverslips were finally washed with PBST three times for 3 min each before embedding on an imaging slide.

Imaging was performed at the Gladstone Institutes Histology and Light Microscopy Core using a Zeiss LSM880 Super-Resolution AiryScan Confocal Microscope.

### Seahorse Respirometry

Prior to experimentation, 96-well Seahorse cell microplates were coated with Cell-Tak (Corning; 354240) per manufacturer instructions. K562 were resuspended in DMEM media (Sigma; D5030) supplemented with 5 mM Hepes, 8 mM Glucose, 2 mM Pyruvate, and 2 mM Glutamine. K562 cells were seeded at 125,000 cells per well and acutely attached by centrifugation. Respiration was measured under basal conditions as well as after injection of 2.5 uM oligomycin (port A), two sequential additions of 0.8 uM FCCP (ports B and C), followed by 0.2 uM rotenone with 1 uM antimycin A (port D). ATP production rate and maximal respiration were calculated as previously described^52^.

### RNAseq

RNA-seq was performed by Plasmidsaurus using Illumina sequencing with a custom library preparation, sequencing, and analysis pipeline. Briefly, 200,000 K562 cells or 300,000 dermal fibroblasts were resuspended in 50µl Zymo DNA/RNA Shield. Poly-A mRNA was captured using an oligo-dT primer incorporating a sample barcode, unique molecular identifier (UMI), and Read 1 sequence, followed by reverse transcription to generate cDNA. Second-strand synthesis produced double-stranded cDNA, which was tagmented to generate fragments and incorporate the Read 2 sequence. Libraries were then amplified with unique dual indices (i5/i7) and P5/P7 adapter sequences for Illumina sequencing.

Quality of the fastq files was assessed using FastQC v0.12.1. Reads were then quality filtered using fastp v0.24.0 with poly-X tail trimming, 3’ quality-based tail trimming, a minimum Phred quality score of 15, and a minimum length requirement of 50 bp. Quality-filtered reads were aligned to the human reference genome (GRCh38, Ensembl release 114) using STAR aligner v2.7.11 with non-canonical splice junction removal and output of unmapped reads, followed by coordinate sorting using samtools v1.22.1. PCR and optical duplicates were removed using UMI-based deduplication with UMIcollapse v1.1.0. Alignment quality metrics, strand specificity, and read distribution across genomic features were assessed using RSeQC v5.0.4 and Qualimap v2.3, with results aggregated into a comprehensive quality control report using MultiQC v1.32. Gene-level expression quantification was performed using featureCounts (subread package v2.1.1) with strand-specific counting, multi-mapping read fractional assignment, exons and three prime UTR as the feature identifiers, and grouped by gene_id. Final gene counts were annotated with gene biotype and other metadata extracted from the reference GTF file. Sample-sample correlations for the sample-sample heatmap and PCA were calculated on normalized counts (TMM, trimmed mean of M-values) using Pearson correlation. Differential expression analysis was performed using edgeR v4.0.16 following standard practice, including filtering of low-expressed genes with edgeR::filterByExpr using default parameters. Functional enrichment, when applicable, was performed using gene set enrichment analysis with gseapy v0.12 against the MSigDB Hallmark gene set collection.

### ChIPseq

#### ChIP-seq library preparation

Chromatin immunoprecipitation (ChIP) was performed in biological replicates as previously described with modifications as follows. For H3K27ac ChIP assay, cells were cross-linked with 1% (vol/vol) formaldehyde in PBS for 10 min at room temperature. For THAP12 and FLAG ChIP assays, cells were cross-linked with 2 mM disuccinimidyl glutarate (DSG) (ProteoChem) in PBS for 30 min at room temperature, and then directly a second crosslinking was performed by the addition of 1% (vol/vol) formaldehyde in PBS for 10 min. The crosslinking reaction was quenched by 0.125 M glycine (Sigma-Aldrich). Cells were washed once with ice-cold PBS and pelleted at 1,000 × g for 5 min at 4°C. Crosslinked cells were resuspended in ice-cold hypotonic buffer (10 mM HEPES-KOH (pH 7.9), 85 mM KCl, 1 mM EDTA, 1% NP-40, 1× protease inhibitor cocktail), and centrifuged at 1,000 × g for 5 min at 4°C to obtain a nuclear fraction. Nuclear pellets were resuspended in 100 μl of either LB3 buffer (10 mM Tris-HCl (pH 7.5), 100 mM NaCl, 1 mM EDTA, 0.5 mM EGTA, 0.1% sodium deoxycholate, 0.5% N-laurosylsarcosine, 1× protease inhibitor cocktail, for H3K27ac ChIP) or PIPA-NR buffer (20 mM Tris-HCl (pH 7.5), 150 mM NaCl, 1 mM EDTA, 0.5 mM EGTA, 0.4% sodium deoxycholate, 0.1% SDS, 1% NP-40, 0.5 mM DTT, 1× protease inhibitor cocktail, for THAP12 and FLAG ChIPs). Chromatin DNA was sonicated in a 0.2 ml microtube (Diagenode) using a Bioruptor® Pico sonication device (30 sec ON / 30 sec OFF cycles) for 15 min at 4°C. Samples were centrifuged at 15,000 rpm for 10 min at 4°C, and the supernatant was used for immunoprecipitation. LB3 lysate was diluted 1.1-fold with 10% Triton X-100. 1% of the lysate was kept as ChIP input. The lysates were rotated with each antibody pre-bound to 20 μl of beads (10 μl of Dynabeads protein A (Thermo Fisher Scientific) + 10 μl of Dynabeads protein G (Thermo Fisher Scientific)) overnight at 4°C (2 μg of H3K27ac antibody (Active Motif; # 39133), 1 μg of THAP12 antibody (BETHYL; #A300-586A), or 2 μg of FLAG antibody (Proteintech; #66008-4-Ig) per assay). After the immunoprecipitation, beads were collected on a magnet and washed three times each with wash buffer I (20 mM Tris-HCl (pH 7.5), 150 mM NaCl, 1% Triton X-100, 0.1% SDS, 2 mM EDTA, 1× protease inhibitor cocktail), wash buffer III (10 mM Tris-HCl (pH 7.5), 250 mM LiCl, 1% Triton X-100, 0.7% sodium deoxycholate, 1 mM EDTA, 1× protease inhibitor cocktail) and twice with TET (10 mM Tris-HCl (pH 7.5), 1 mM EDTA, 0.2% Tween-20, 1× protease inhibitor cocktail), and then resuspended in 25 μl of TT (10mM Tris-HCl (pH 7.5), 0.05% Tween-20). The immunoprecipitated chromatin samples were used for library preparation with NEBNext Ultra II Library kit (NEB) according to the manufacturer’s instructions. DNA in 46.5 μl of NEB reactions was reverse crosslinked by adding 4 μl of 10% SDS, 4.5 μl of 5 M NaCl, 3 μl of 500 mM EDTA, 1 μl of 20 mg/ml proteinase K and 20 μl of water by incubation for 1 hr at 55°C, and then 30 min at 65°C. DNA was purified with 2 μl of SpeedBeads in 20% PEG 8000/1.5 M NaCl (Final 12% PEG 8000), and eluted with 25 μl of TT. The eluted DNA was amplified for 14 cycles in 50 μl of PCR reactions using NEBNext High-Fidelity 2× PCR Master Mix (NEB) and 0.5 mM each of primers Solexa 1GA and Solexa 1GB. The amplified libraries were purified with 2 μl of SpeedBeads in 20% PEG 8000/2.5 M NaCl (Final 13% PEG 8000), eluted with 20 μl of TT, size selected using PAGE/TBE gel for 225-325 bp fragments by gel extraction, and single-end sequenced on NovaSeq X Plus (Illumina). ChIP input material (1% of sheared DNA) in 46.5 μl of TE was reverse crosslinked by adding 4 μl of 10% SDS, 4.5 μl of 5 M NaCl, 3 μl of 500 mM EDTA, 1 μl of 20 mg/ml proteinase K and 20 μl of water by incubation for 1 hr at 55°C, and then 30 min at 65°C. The input DNA was purified with SpeedBeads as described above, and eluted with 25 μl of TT. The eluted input DNA was prepared for libraries and amplified as described for ChIP samples.

#### ChIP-seq data analysis

FASTQ files were mapped to the hg38 human reference genome with Bowtie2. Peaks were called with HOMER (findPeaks) using parameters “-style factor -minDist 200 -size 200”. After merging these peaks, correlations among replicates from the same cell subset/treatment were evaluated by correlation using tag counts. The two most highly correlated samples were used for identifying the most robust peaks using the irreproducible discovery rate (IDR) method. For this step, peaks were called with HOMER’s findPeaks, using parameters “-L 0 -C 0 -fdr 0.9 -minDist 200 -size 200”. IDR peaks from different conditions involved in a comparison merged with HOMER’s mergePeaks and annotated with HOMER’s annotatePeaks.pl. The raw tags of all samples which had reasonable correlation were quantified with HOMER (annotatePeaks.pl) using parameter “-noadj”. Peaks which contained at least 4 tags in at least 1 sample were used to identify differentially bounded peaks (DBP) by DESeq2. Peaks were categorized as distal peaks which are 2 kb away from known TSS and promoter peaks which are located within the 2 kb region of known TSS sites. ChIP-seq data were visualized in the UCSC genome browser.

## ACKNOWLEDGEMENTS/FUNDING

We thank all present and past members of the Jain Lab, as well Andrea Curtabbi for their thoughtful discussion and review of the manuscript. We greatly thank Vinh Nguyen from the UCSF Parnassus Flow CoLab for assistance with CRISPR screen FACS sorting. We give our deepest thanks to Jane Srivastava from the Gladstone Institutes Flow Cytometry Core and Rebecca Blandino from the Histology and Light Microscopy Core. We thank Dr. Gabriela Grigorean at UC-Davis for assistance with proteomics. We thank Rebeca Acín-Pérez for technical support with BN-PAGE. We thank Dr. Françoise Chanut and Dr. Brian Plosky for their editorial support during the writing of this work. We thank Tami Tolpa for her assistance with data visualization and graphics.

CRISPR screen sequencing was performed at the UCSF CAT, supported by UCSF PBBR, RRP IMIA, and NIH 1S10OD028511-01 grants. Funding was provided by the Department of Defense National Defense Science and Engineering Graduate Fellowship (NDSEG, B.R.D.), NIH DP5 DP5OD026398 (I.H.J.), Klingenstein-Simons Fellowship (I.H.J.).

## ETHICS DECLARATIONS

Competing interests: I.H.J. has patents related to hypoxia-based therapies.

## SUPPLEMENTAL FIGURES

**Supplemental Figure 1.**
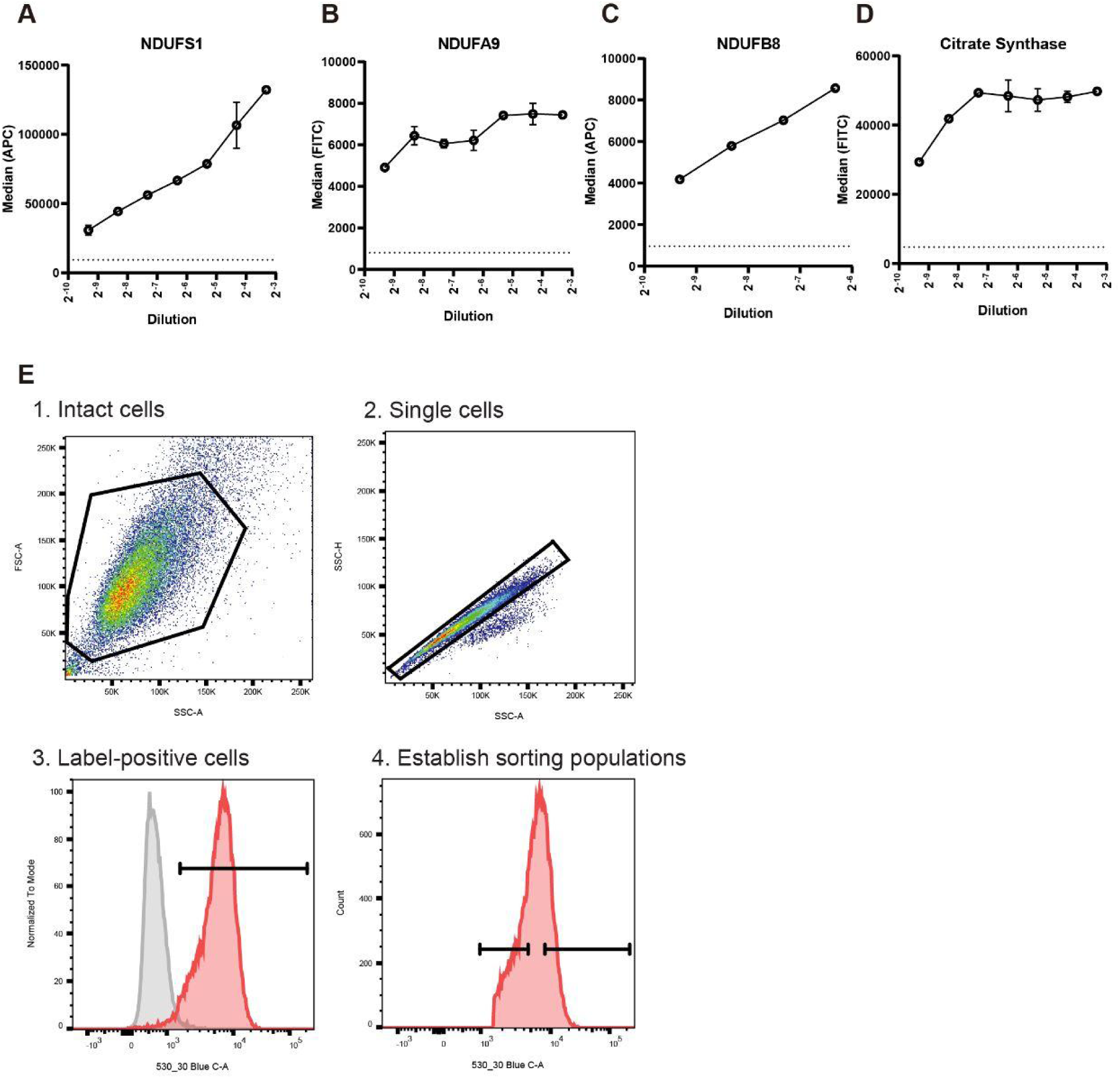
Optimization of primary antibody staining and depiction of gating strategy for FACS CRISPR screen. Primary antibody titration curves depicted for antibodies used in screens. Antibodies include NDUFS1 (A), NDUFA9 (B), NDUFB8 (C, and Citrate Synthase (D). (B) Depiction of gating strategy used for FACS screen. Samples were first gated for intact (1) and singularized (2) cells based on their forward- and side light scattering. Singularized cells are assessed for fluorescence signal by comparing against a secondary antibody-only negative control (3). Cells with true positive fluorescence were then gated based on their relative fluorescence signal. Cells within the bottom 30% or top 30% for fluorescence signal were collected for barcode sequencing.

**Supplemental Figure 2.**
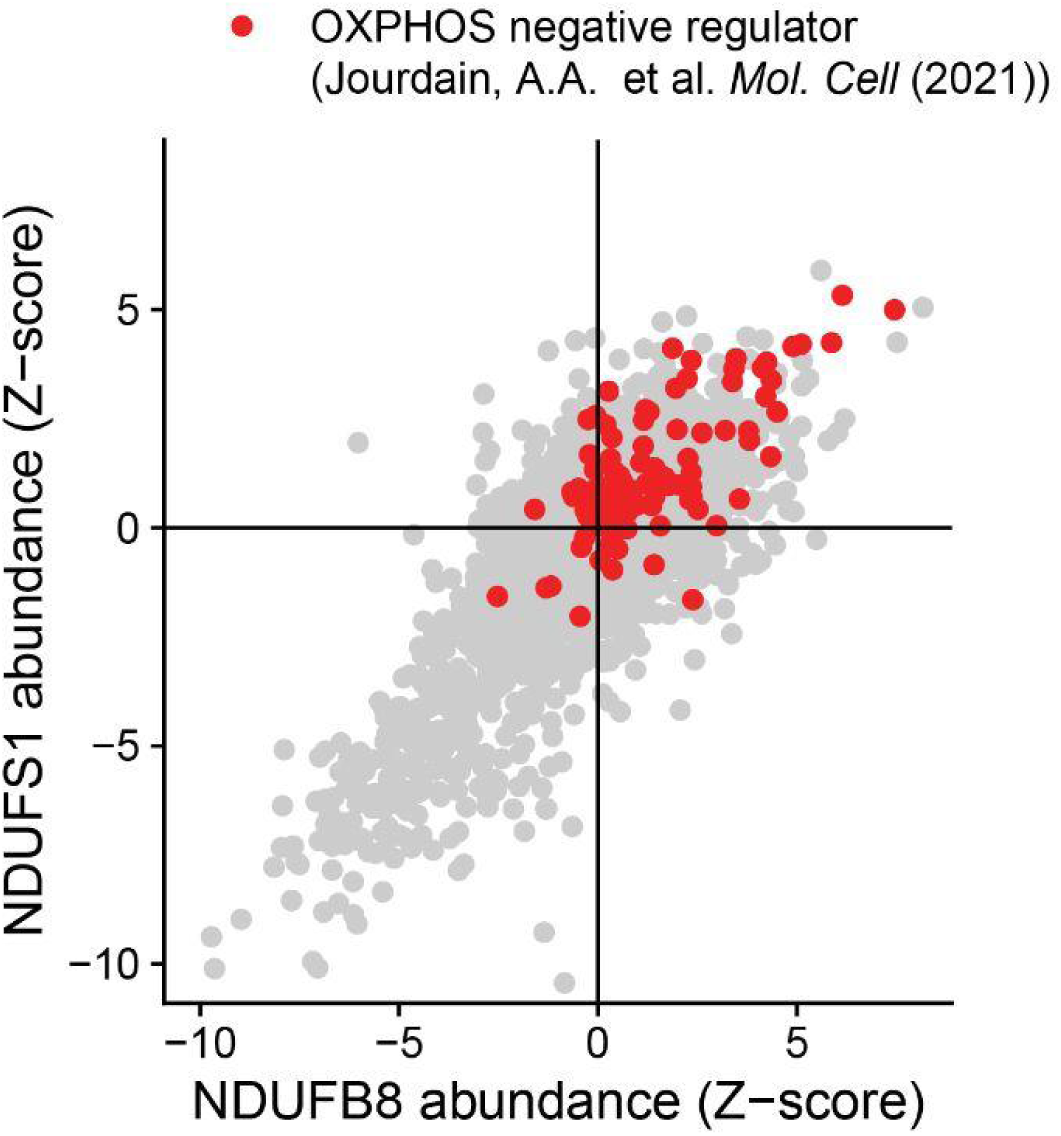
Comparison of Complex I screen results to galactose viability screen. NDUFS1 and NDUFB8 CRISPR screen compared against each. Top t0 hits from a previously performed galactose are denoted in red.

**Supplemental Figure 3.**
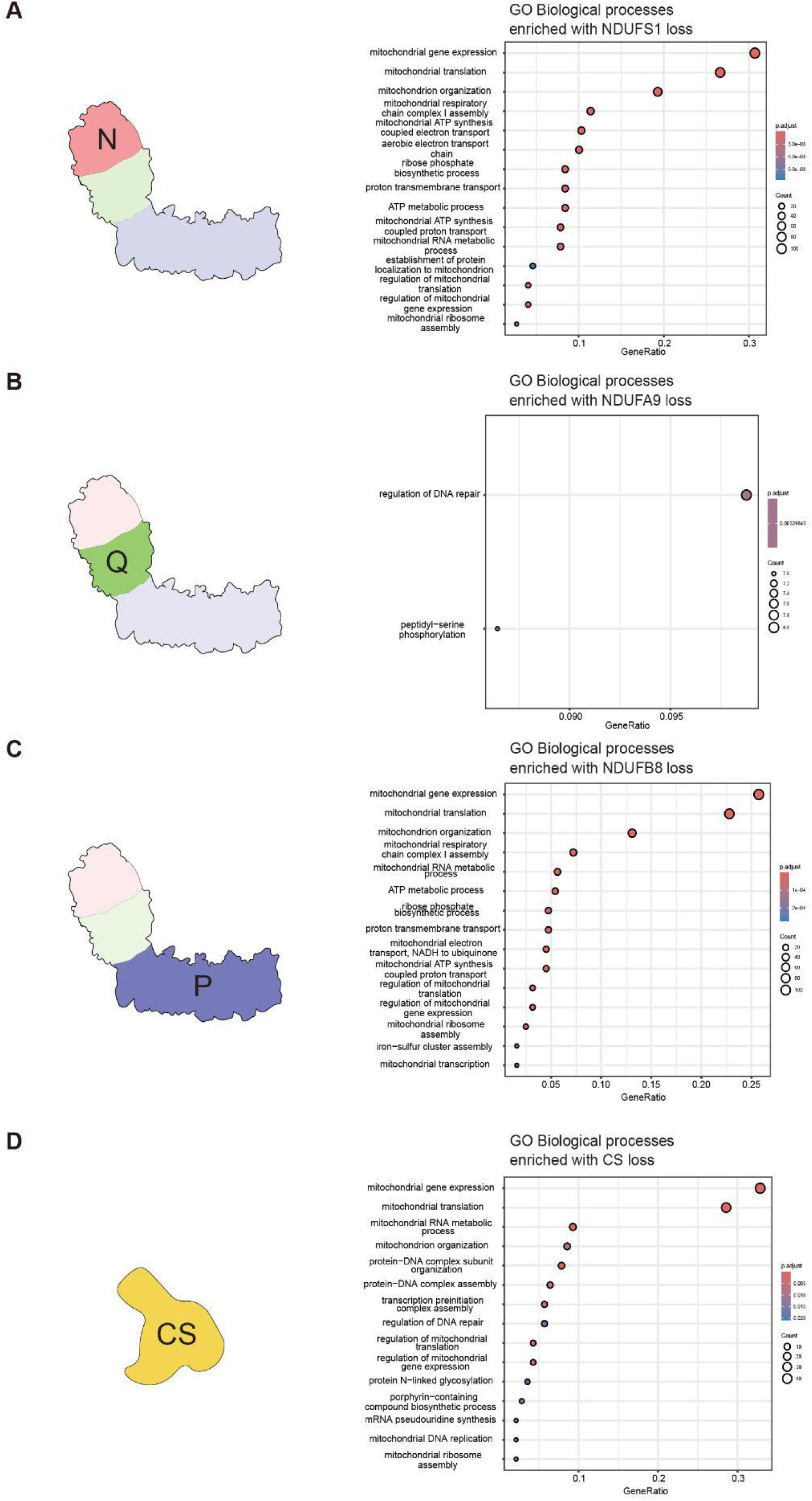
Summary of biological processes associated with Complex I loss across CRISPR screens. Gene ontology assessment for biological processes represented on the depleting side of the screens. Biological processes are depicted for the NDUFS1 (A), NDUFA9 (B), NDUFB8 (C, and Citrate Synthase (D) screens.

**Supplemental Figure 4.**
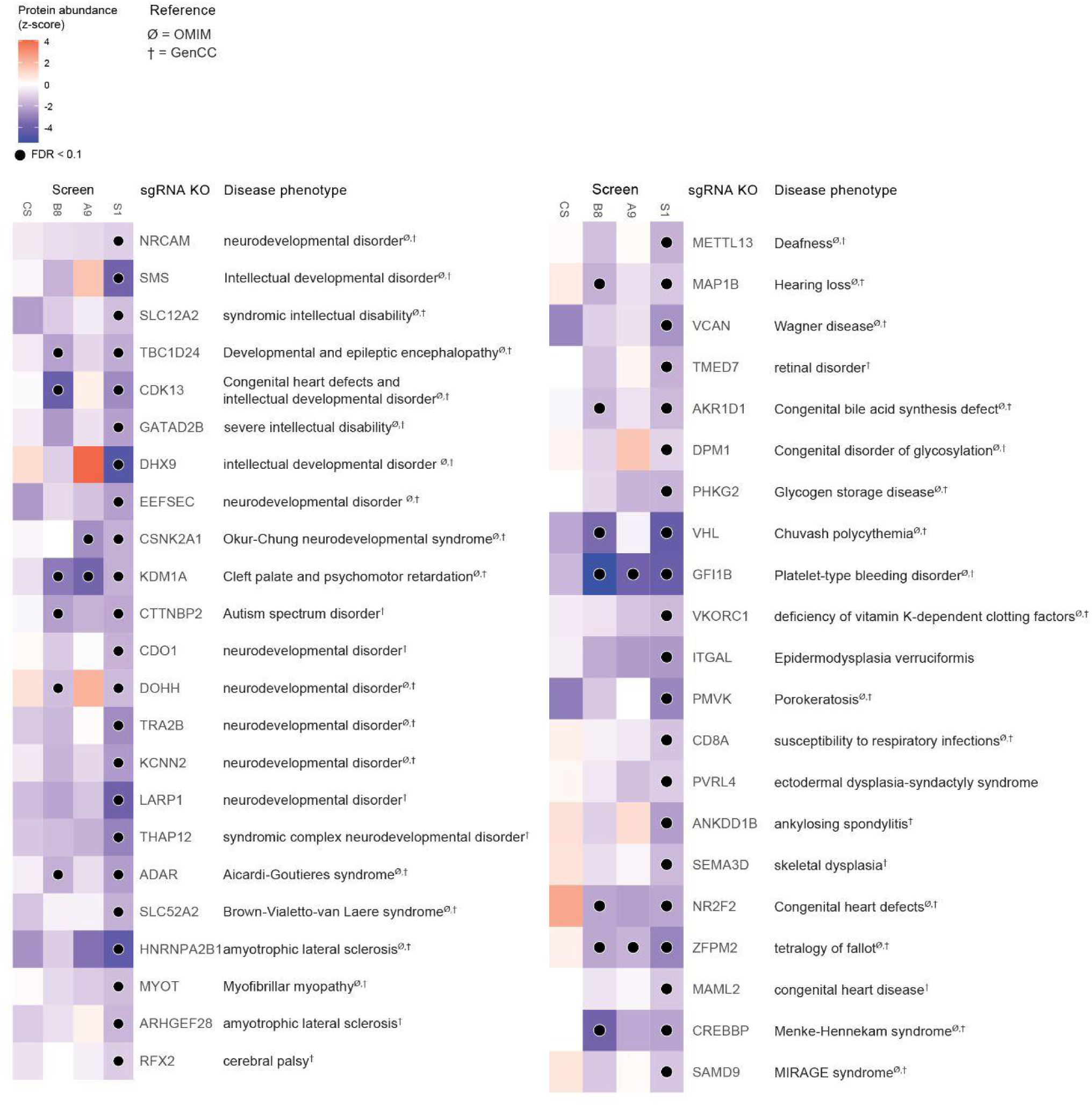
Human disease phenotypes associated with non-mitochondrial modifiers of Complex. **I.** Non-mitochondrial genes, as evaluated by their presence in MitoCarta 3.0, were filtered based on the presence of a human disease phenotype. Human disease phenotypes were identified based on their presence in Online Mendelian Inheritance in Man (OMIM) and Gene Curation Coalition (GenCC) databases.

**Supplemental Figure 5.**
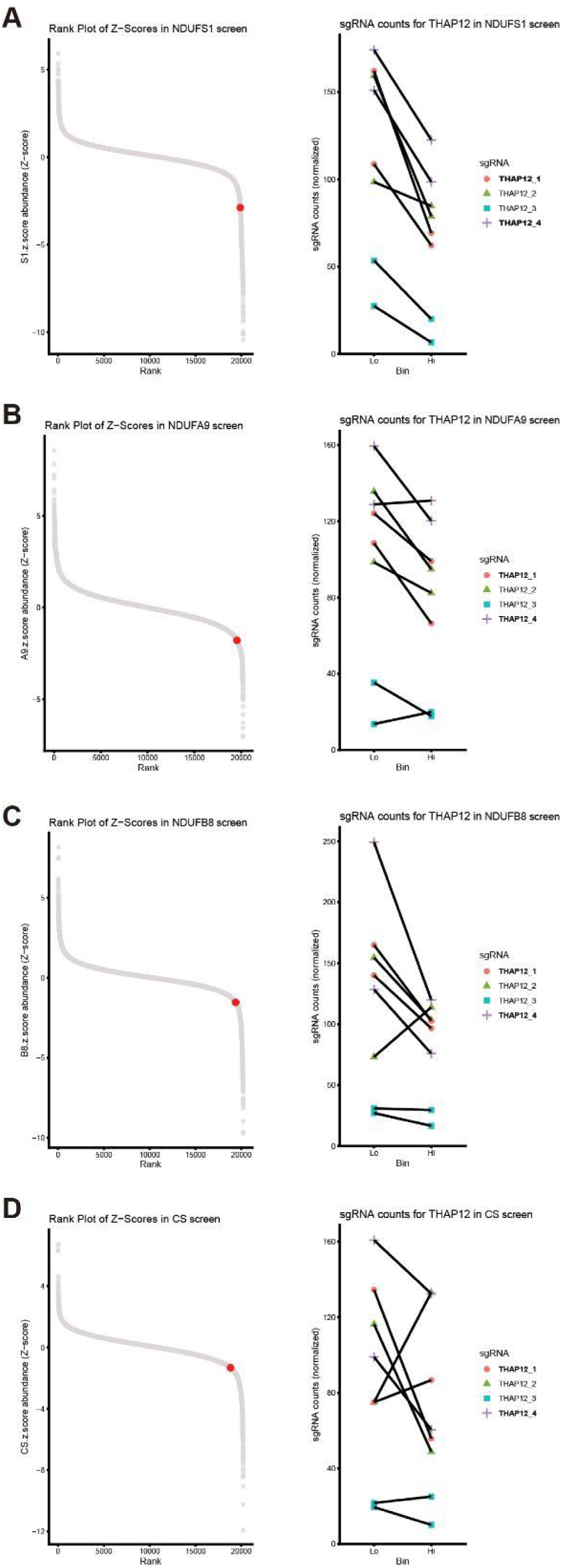
THAP12 results by screen. THAP12 results from the NDUFS1 (A), NDUFA9 (B), NDUFB8 (C), and Citrate Synthase (D) screens. For each screen, THAP12 is highlighted in the rank plot (left) alongside its per-guide results (right).

**Supplemental Figure 6.**
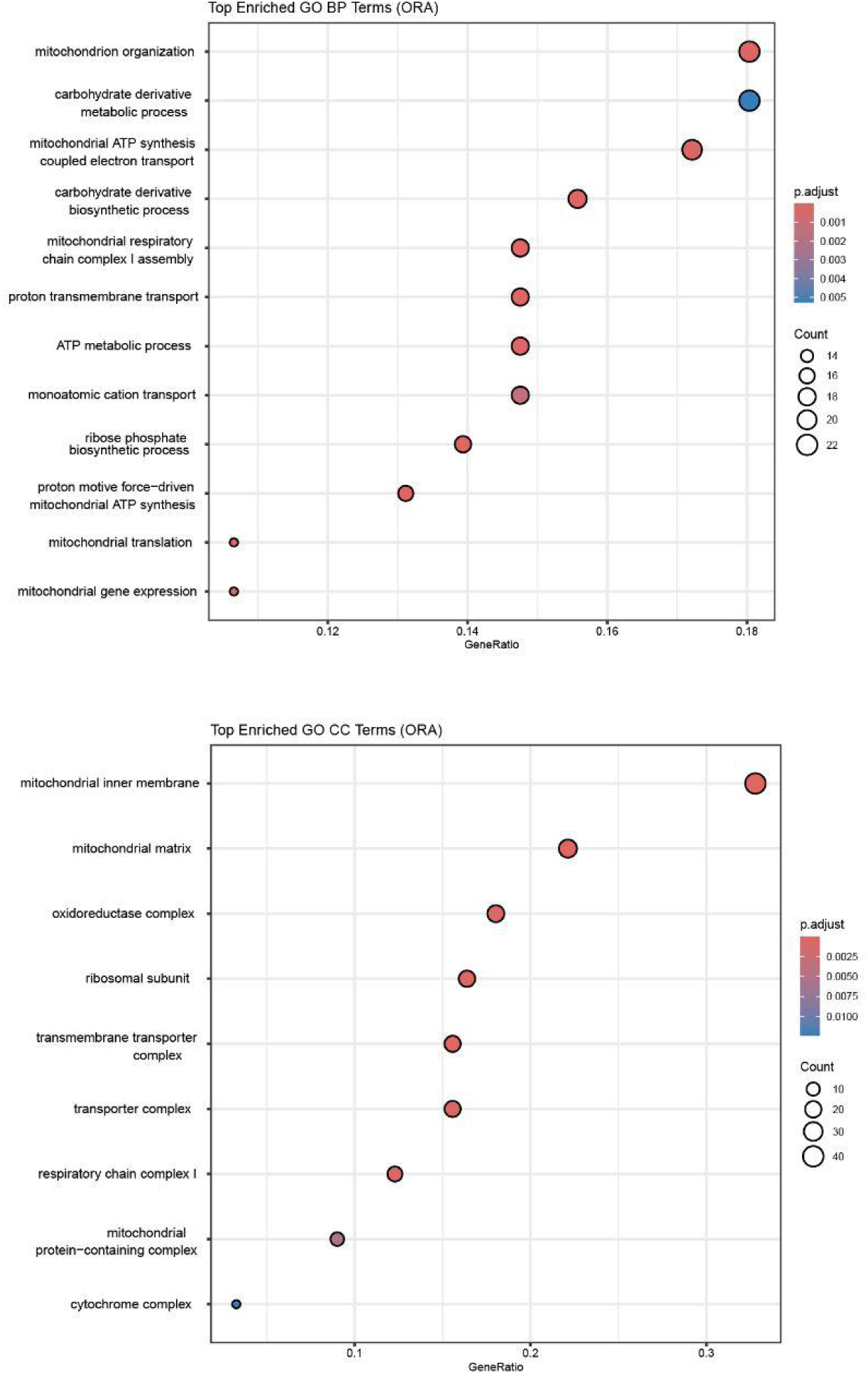
Disrupted processes identified in THAP12 KO proteomics. Gene ontology analysis of proteomics data from THAP12 KO K562 cells depicting the biological processes (top) and cellular compartments (bottom) associated with the most depleted proteins.

**Supplemental Figure 7.**
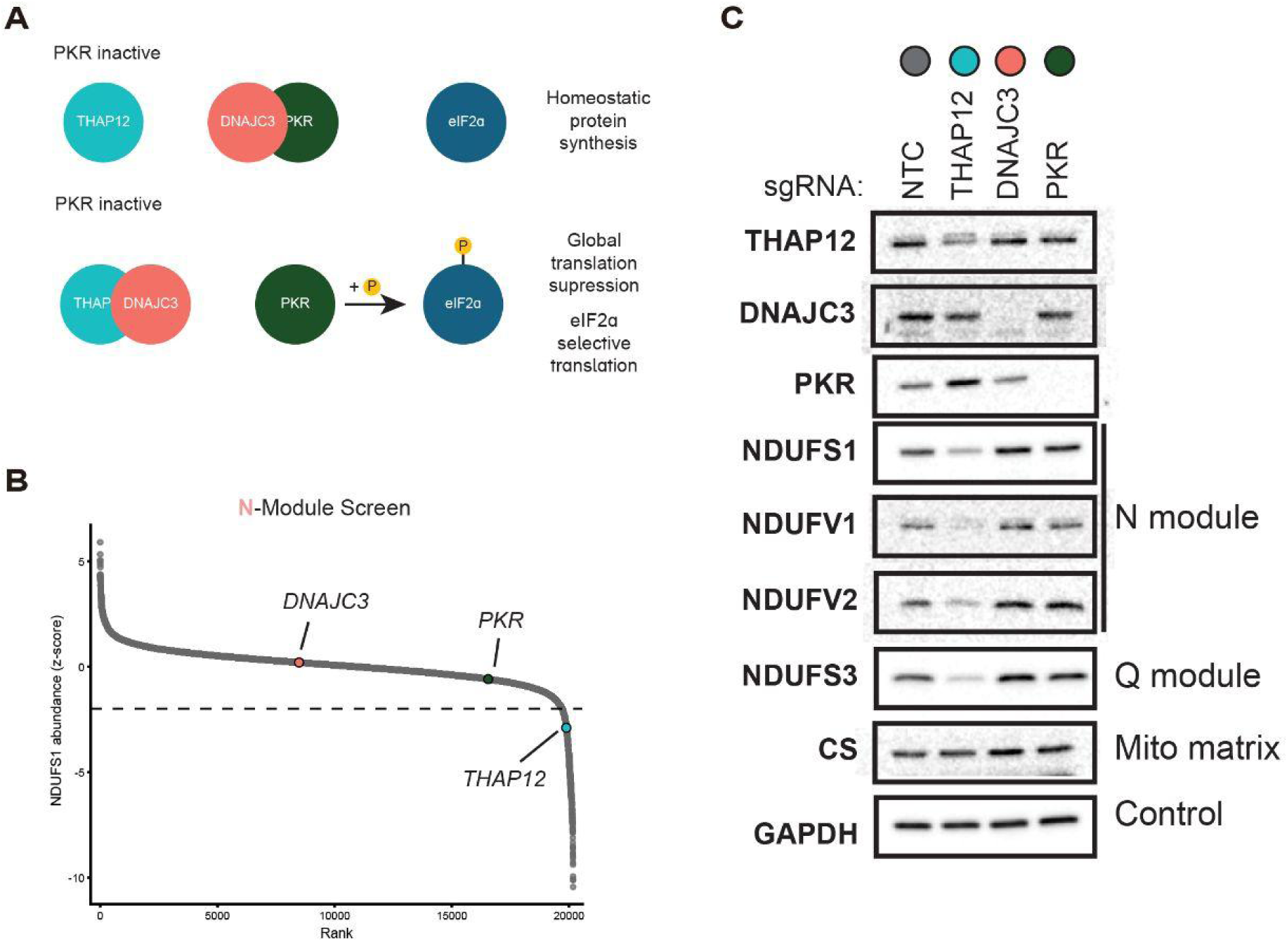
Disrupted PKR signaling does not explain CI deficiency with THAP12 loss. (A) Previously proposed model for THAP12 in regulating PKR signaling through direct interaction with DNAJC3. (B) Evaluation of results for PKR signaling members in the NDUFS1 N module screen. (C) Assessment of CI protein abundance with sgRNA KOs of individual members of the PKR signaling cascade.

**Supplemental Figure 8.**
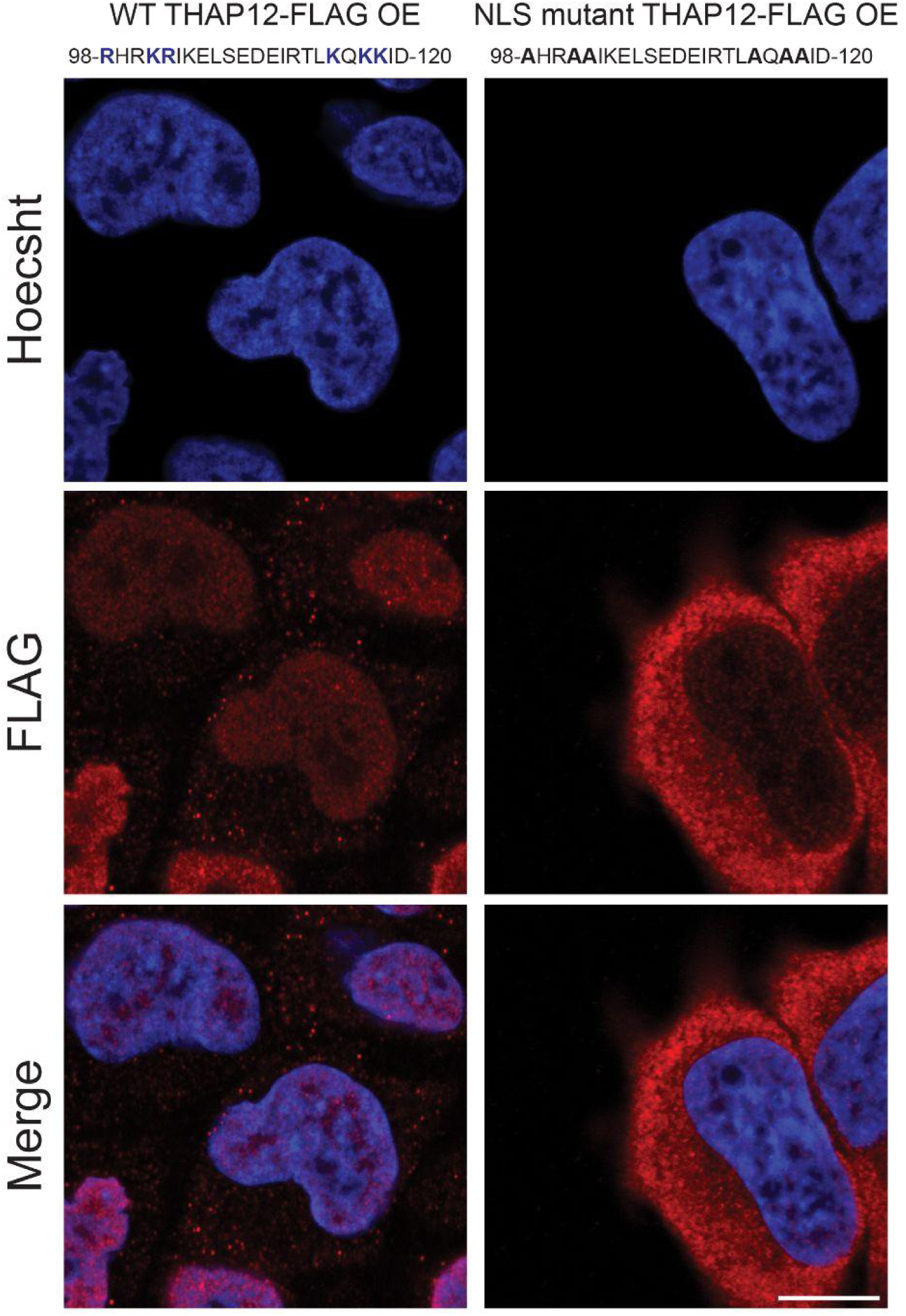
THAP12 nuclear localization is largely coordinated by a bipartite nuclear localization signal (NLS). Assessment of THAP12 localization after neutralizing a predicted NLS sequence. Basic residues imparting a positive charge for nuclear localization are depicted in blue (left). These basic residues were converted to alanine to neutralize the positive charge. These changes are depicted with bold, black lettering (right).

**Supplemental Figure 9.**
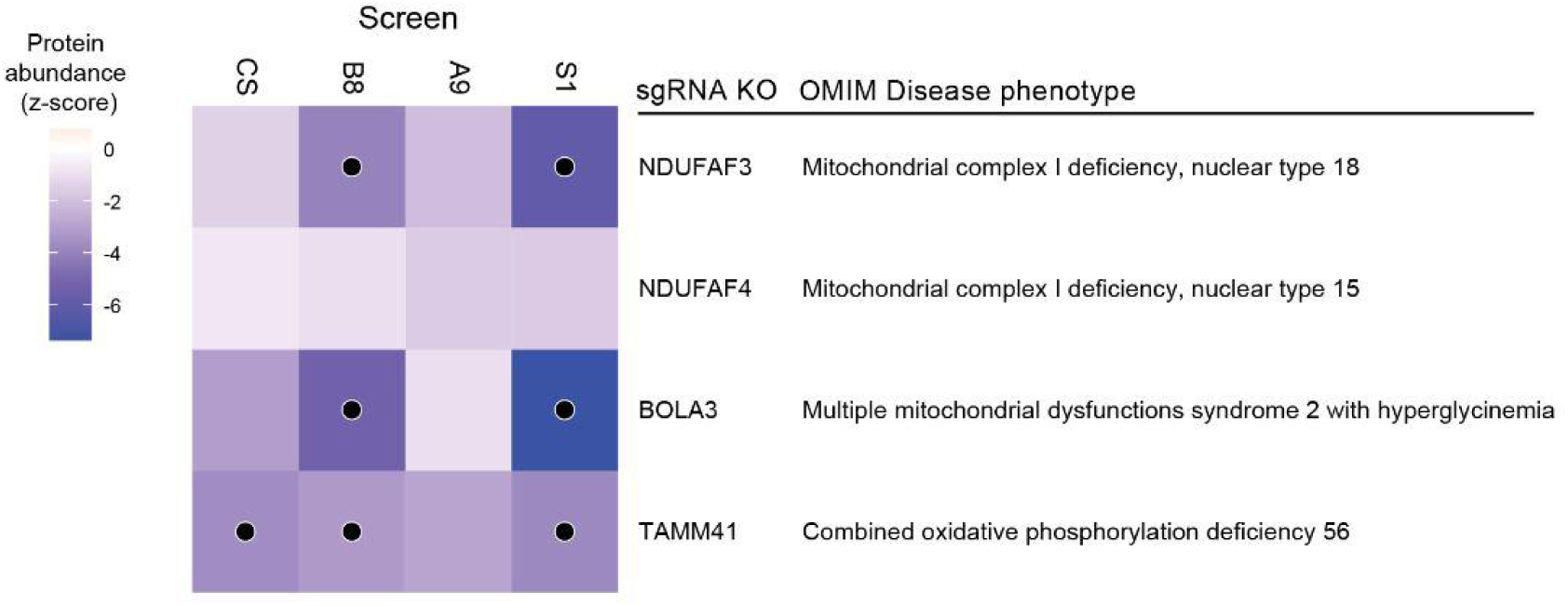
THAP12 controls a transcriptional node required for CI biogenesis and implicated in mitochondrial disease. Results from CI CRISPR screens for defined THAP12 gene targets. Cells are labeled with a dot to denote an FDR < 0.1. THAP12 targets are additionally annotated with their respective disease phenotype annotation present in OMIM.

**Supplemental Figure 10.**
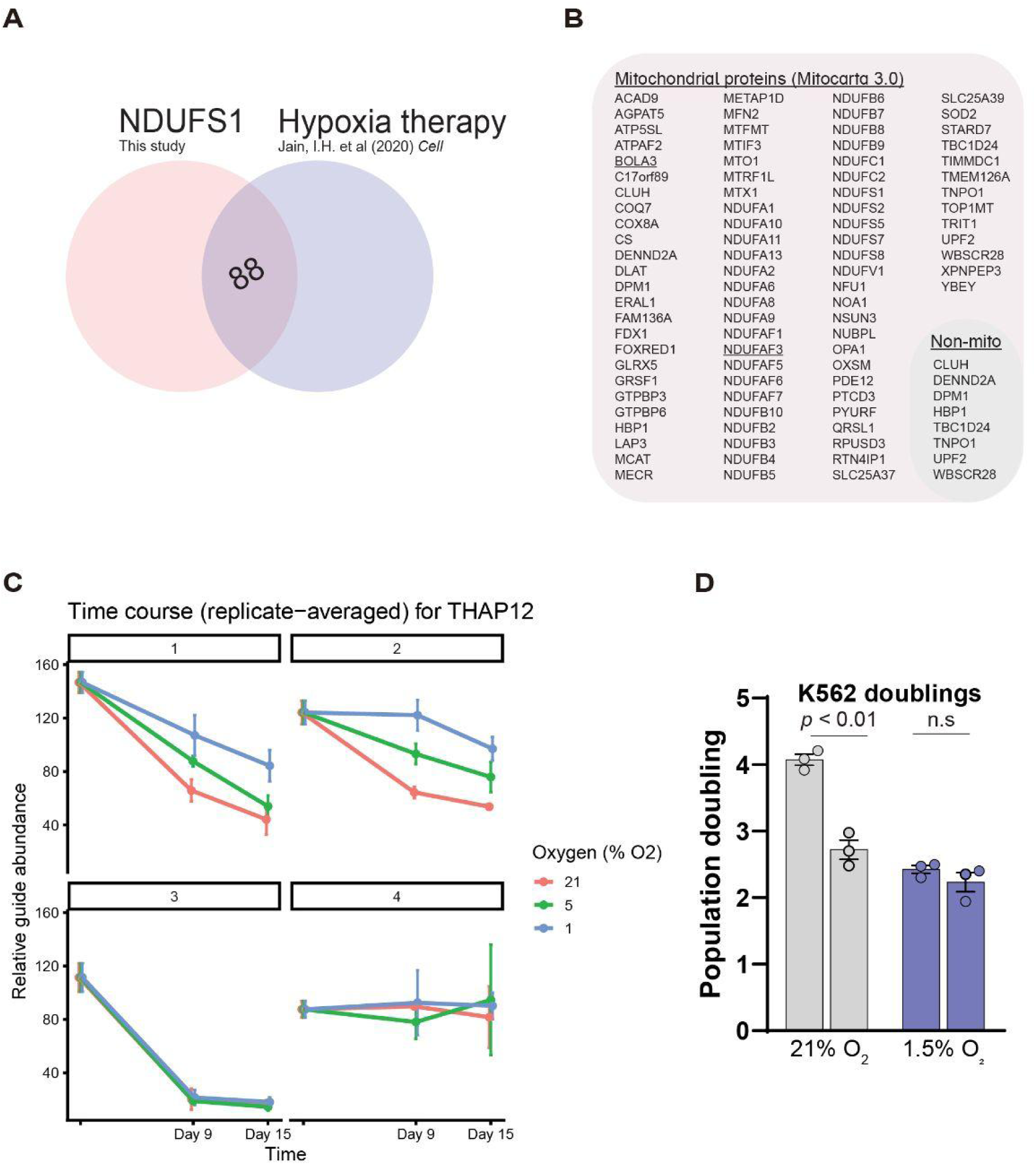
THAP12 disease as a candidate for hypoxia therapy. (A) Comparison of screen results from the NDUFS1 screen (this work) and the previously performed hypoxia therapy screen (B) List of genes identified in screen comparison analysis that are further grouped based on mitochondrial localization. (C) Guide-level analysis for THAP12 sgRNA KO cells in the hypoxia therapy screen, relevant to Fig. 6E. (D) Validation of hypoxia therapy result using monoclonal THAP12 KO K562 cells. This data is a non-normalized representation of data presented in Fig. 6F

## SUPPLEMENTARY TABLE LEGENDS

**Supplemental Table 1. CRISPR screen MAGeCK results.** MAGeCK analysis output for the FACS-based Complex I screens performed. Gene-level statistics data from each CRISPR screen is presented on their own individual sheet.

**Supplemental Table 2. Whole cell proteomics analysis**. Whole cell proteomics analysis generated using the limma analysis package. Proteomics for THAP12 KO and NDUFS1 KO cells are presented on individual sheets.

**Supplemental Table 3. RNAseq differential gene expression analysis.** Differential gene expression analysis from THAP12 KO K562 cells and THAP12 mutant patient fibroblasts. K562 and fibroblast data are presented on their own individual sheets.

